# Tissue-specific modifier alleles determine *Mertk* loss-of-function traits

**DOI:** 10.1101/2022.05.23.493101

**Authors:** Yemsratch T. Akalu, Maria E. Mercau, Marleen Ansems, Sagie Wagage, Lindsey D. Hughes, James Nevin, Emily Alberto, Xinran Liu, Li-Zhen He, Diego Alvarado, Tibor Keler, Yong Kong, William M. Philbrick, Silvia C. Finnemann, Antonio Iavarone, Anna Lasorella, Carla V. Rothlin, Sourav Ghosh

## Abstract

Knockout (KO) mouse models play critical roles in elucidating biological processes behind disease-associated or disease-resistant traits. As a consequence of gene KO, mice display certain phenotypes. Based on insight into the molecular role of said gene in a biological process, it is inferred that the particular biological process causally underlies the trait. This approach has been crucial towards understanding the basis of pathological and/or advantageous traits associated with *Mertk* KO. MERTK is a receptor tyrosine kinase with a critical role in phagocytosis of apoptotic cells or cellular debris. Therefore, early-onset, severe retinal degeneration was described to be a direct consequence of failed phagocytosis of photoreceptor outer segments by retinal pigment epithelia. Similarly, enhanced anti-tumor immunity was inferred to result from the failure of macrophages to dispose cancer cell corpses, resulting in a pro-inflammatory tumor microenvironment. Here we report that the loss of *Mertk* alone is not sufficient for retinal degeneration. This trait only manifests when the function of the paralog *Tyro3* is concomitantly lost. Additionally, the dramatic resistance against two syngeneic mouse tumor models observed in *Mertk* KO cannot, at least entirely, be ascribed to the loss of *Mertk*. The widely used *Mertk* KO carries multiple coincidental changes in its genome that affect the expression of a number of genes, including *Tyro3*. Nonetheless, neither *Tyro3*, nor macrophage phagocytosis by alternate genetic redundancy, accounts for the absence of anti-tumor immunity in two independent *Mertk* KOs. Collectively, our results indicate that context-dependent epistasis of independent modifier alleles determine *Mertk* KO traits.

## INTRODUCTION

The receptor tyrosine kinase (RTK) MERTK is a paralog of TYRO3 and AXL and together these receptors are commonly referred to as TAM RTKs. *Mertk* was named after its expression pattern in <u>m</u>onocytes, <u>e</u>pithelial tissues and reproductive tissues and for it being a tyrosine <u>k</u>inase^1^. An understanding of MERTK’s role in molecular and cellular processes, as well as its broader role in mammalian physiology and pathology came, in large part, from the generation of a *Mertk* ^-/-^ mouse line established by Camenisch *et al* ^2^. Use of this *Mertk* ^-/-^ mouse line revealed the critical functional role of this RTK in downregulation of inflammatory cytokines such as TNFα, as well as in the phagocytosis and clearance of apoptotic thymocytes ^2,3^. Subsequently, the *Mertk* ^-/-^ mouse line became the fountainhead for the description of *Mertk* function in a spectrum of phenotypes spanning retinal degeneration, defective adult neurogenesis, neurodegenerative diseases, liver injury, lupus-like autoimmunity and cancer ^4-15^.

The *Mertk* ^-/-^ mouse line was generated by using the available technology of the time. Specifically, *Mertk* was targeted in 129P2/OlaHsd (129P2)-derived E14TG2a embryonic stem (ES) cells ^2^. ES cells were then microinjected into a C57BL/6 (B6) blastocyst to generate a chimeric mouse with germline transmission of the targeted allele (**Fig. 1A, B**). Subsequently, the chimeric mouse was backcrossed to B6 to obtain *Mertk* ^-/-^ mice, henceforth referred to as *Mertk* ^-/- V1^ (**Fig. 1A, Fig S1A, B**). The *Mertk* ^-/- V1^ mouse line is available through The Jackson Laboratory (Strain #: 011122). It is typically backcrossed >10 generations into B6 mice by researchers, including us, and has remained the mainstay for MERTK research. Nevertheless, there have been occasional and isolated reports of independently generated *Mertk* knockout mice that failed to completely recapitulate *Mertk* ^-/- V1^ phenotypes ^16^. For example, early-onset, severe photoreceptor (PR) degeneration was reported in *Mertk* ^-/- V1^ mice ^4,17,18^. In these mice, the outer nuclear layer (ONL) thickness was significantly reduced by postnatal day (P) 25 ^4^. Electroretinogram (ERG) recordings revealed that scotopic a- and b-wave amplitudes were significantly lower in *Mertk* ^-/- V1^ mice at P20 when compared to wild type (WT) mice at P30 ^4^. Photopic amplitudes were also significantly lower in *Mertk* ^-/- V1^ mice *versus* WT mice at P33 ^4^. In a different study, an independently generated ENU-induced *Mertk* mutation (*Mertk* ^nmf12 or H716R^) in B6 mice caused the substitution of a highly conserved histidine to an arginine and led to a drastic reduction of MERTK in mouse retinas ^16^. Yet, it did not identically phenocopy the *Mertk* ^-/- V1^-associated early-onset, severe retinal degeneration. Since a slow form of retinal degeneration did indeed occur in *Mertk* ^nmf12 or H716R^ and MERTK expression was not entirely abolished ^16^, potential problems with *Mertk* ^-/- V1^ mice were not immediately brought to the fore.

**Figure 1.**
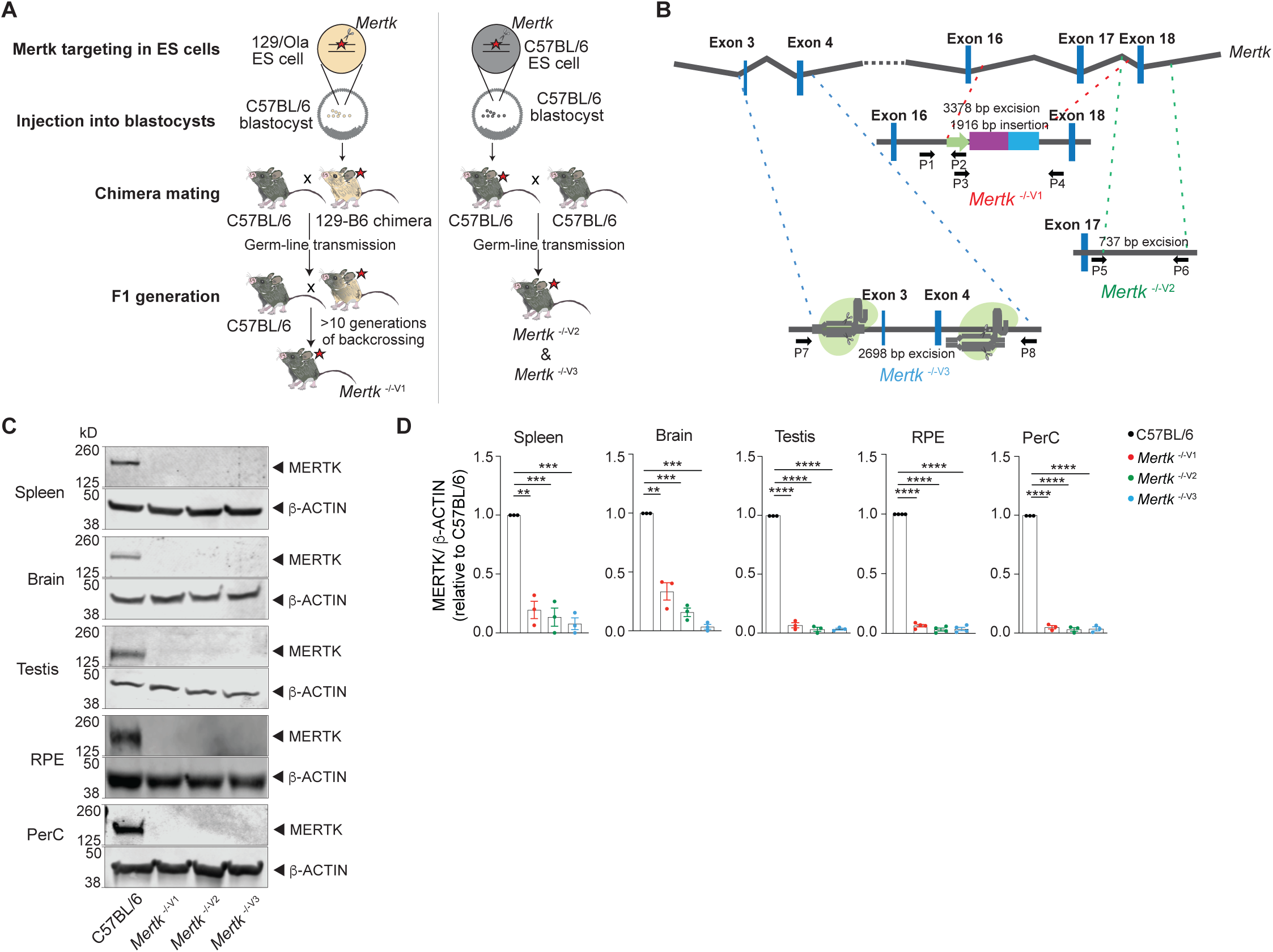
Generation of B6 embryonic stem (ES) cell-derived mice with genetic ablation of *Mertk*. (**A**) Schematic showing the differences in approach between the generation of *Mertk* ^-/- V1^ mice by targeting *Mertk* in 129P2/OLAHsd (129/Ola)-derived ES cells by Camenisch *et al*. and our *Mertk* knockout mouse lines. 129P2 ES cells were microinjected into C57BL/6 (B6) blastocysts to generate chimeric mice with germline transmission of deleted *Mertk* allele by Camenisch *et al*. Chimeric mice were subsequently backcrossed onto B6 mice, for >10 generations in our laboratory to obtain *Mertk* ^-/- V1^ mice. Our two independent *Mertk* knockout mouse lines, *Mertk* ^-/- V2^ and *Mertk* ^-/- V3^ mice, were generated by targeting *Mertk* in B6 ES cells. Red stars indicate at least one copy of the mutant allele of *Mertk*. (**B**) Schematic indicating *Mertk* ^-/- V1^ mice have deletion of exon 17 that encodes for the kinase domain of *Mertk*. A neomycin cassette is also present at this site. *Mertk* ^-/- V2^ mice have targeted excision of exon 18 which also encodes for residues in the kinase domain. *Mertk* ^-/- V3^ mice have exons 3 and 4 targeted with CRISPR/Cas9 approach. (**C, D**) Representative and quantification of independent MERTK western blot data depicting total MERTK protein expression in spleen, brain, testes, retinal pigment epithelia (RPE) and peritoneal cavity cells (PerC) from C57BL/6, *Mertk* ^-/- V1^, *Mertk* ^-/- V2^ and *Mertk* ^-/- V3^ mice. (mean ± SEM of n=3-4 mice/genotype). **p<0.01. ***p<0.001, ****p<0.0001, one-way ANOVA-Dunnet’s test.

In an independent study, Vollrath *et al*. demonstrated that crossing *Mertk* ^-/- V1^ mice to B6 mice occasionally gave rise to animals with normal retina ^19^. The authors further demonstrated that the *Mertk* ^-/- V1^ mice carry a ∼40 cM segment around *Mertk* derived from 129P2 strain. The very low frequency of normal retina phenotype indicates that crossovers are extremely rare within this chromosomal segment around *Mertk*. After rare crossover events, when B6 alleles within this region were present, *Mertk* ^-/- V1^-dependent retinal degeneration was prevented ^19^. Vollrath *et al*. mapped the suppressor of retinal degeneration to a region that encoded 53 known or predicted open reading frames (ORFs). *Tyro3* was identified *a priori* as the likely candidate providing the suppressor function since it is a paralog of *Mertk*. Consistent with this hypothesis, it was observed that TYRO3 expression was at ∼33% for *Tyro3* ^129/129^ (*e.g*. in *Mertk* ^-/- V1^) relative to *Tyro3* ^B6/B6^ amounts, and associated with retinal degeneration ^19^. By contrast, TYRO3 expression was at ∼67% for *Tyro3* ^B6/129^ relative to *Tyro3* ^B6/B6^ amounts. This ∼67% expression of TYRO3 prevented retinal degeneration.

These results indicate that not all phenotypes in *Mertk* ^-/- V1^ mice are solely due to the loss of MERTK function. Such crucial anomalies notwithstanding, this *Mertk* ^-/- V1^ mouse line continues to be used to ascribe pivotal functions to MERTK in wide ranging diseases such as neurodegeneration and cancer. Therefore, we investigated if indeed phenotypes observed in *Mertk* ^-/- V1^ mice can be solely and unambiguously ascribed to the loss-of-function of *Mertk*. Here we show that two independently generated B6 *Mertk* ^-/-^ mice do not phenocopy two of the major phenotypes – retinal degeneration and anti-tumor resistance – characteristic of *Mertk* ^-/- V1^ mice. Furthermore, complementary to genetic evidence that retinal degeneration segregated with *Tyro3* ^129/129^ but not *Tyro3* ^129/B6^ reported by Vollrath *et al*., we demonstrate that the simultaneous ablation of *Mertk* and *Tyro3* in B6 mice is necessary and sufficient for retinal degeneration. MERTK and TYRO3 share the two most well-described TAM functions – phagocytosis and anti-inflammatory signaling ^20^. Thus, functional redundancy provided by TYRO3 in the phagocytosis of photoreceptor outer segments (POS) by retinal pigment epithelia (RPE) in the absence of MERTK is consistent with the well-understood mechanism of retinal homeostasis. Nevertheless, loss of MERTK is also proposed to hinder macrophage-dependent phagocytosis of dead or dying cancer cells (efferocytosis). This deficiency in macrophage-mediated disposal of tumor cells, in turn, is postulated to improve availability of tumor antigens for proficient presentation on dendritic cells (DCs), and/or render the tumor microenvironment pro-inflammatory and less immunosuppressive ^11,12,21^. Paradoxical to this view, macrophages from *Mertk* ^-/- V1^, *Mertk* ^-/- V2^ or *Mertk* ^-/- V3^ mice all displayed significant deficit in efferocytosis. Even mice with simultaneous ablation of *Mertk* and *Tyro3* did not phenocopy the anti-tumor resistance of *Mertk* ^-/- V1^ mice. RNA sequencing of bone marrow-derived macrophages (BMDMs) and RPE revealed changes in expression of ∼12- 16 genes located in chromosome 2 in *Mertk* ^-/- V1^ but not *Mertk* ^-/- V2^ or *Mertk* ^-/- V3^ mice. Changes in the expression of additional non-linked genes beyond chromosome 2 were also observed in *Mertk* ^-/- V1^ but not *Mertk* ^-/- V2^ or *Mertk* ^-/- V3^ mice. Furthermore, this differential gene expression between *Mertk* ^-/- V1^ and *Mertk* ^-/- V2^ or *Mertk* ^-/- V3^ mice were tissue-specific, pointing to the presence of a number of modifier alleles that may function combinatorially in several cell-types for at least some of the *Mertk* ^-/- V1^ mouse traits.

## RESULTS

### *Mertk* ablation in B6 ES cells is not sufficient to cause retinal degeneration

We engineered two new *Mertk* knock out mouse lines (designated *Mertk* ^-/- V2^ and *Mertk* ^-/- V3^ mice) generated directly using B6 ES cells **(Fig. 1A, B)**. In the first strategy, we ablated exon 18 within the region encoding the kinase domain and containing the critical ATP-coordinating lysine residue (*Mertk* ^-/- V2^ mice; **Fig. 1B, Fig. S1B**). In an independent approach, we employed CRISPR/CAS9 to delete exons 3 and 4 of *Mertk* (*Mertk* ^-/- V3^; **Fig. 1B, Fig. S1C**). Immunoblotting of lysates from a variety of tissues, including the spleen, brain, testis, cells from the RPE and peritoneal cavity (PerC), validated that no detectable MERTK was observed in *Mertk* ^-/- V2^ and *Mertk* ^-/- V3^ mice (**Fig. 1C, D**). The reduction in MERTK amounts in *Mertk* ^-/- V2^ and *Mertk* ^-/- V3^ mice was comparable to that in *Mertk* ^-/- V1^ tissues (**Fig. 1C, D**).

Next, we investigated whether retinal degeneration characteristic of the *Mertk* ^-/- V1^ mouse is phenocopied in *Mertk* ^-/- V2^ and *Mertk* ^-/- V3^ mice. We performed histological analyses as well as transmission electron microscopy of retinal sections at 6-months of age. As expected, *Mertk* ^-/- V1^ mice displayed advanced PR loss, evidenced by the presence of ∼1 row of nuclei in the ONL across the entire dorsal-ventral axis of the retina (**Fig. 2A, B**). By contrast, morphological analysis of retinas from *Mertk* ^-/- V2^ and *Mertk* ^-/- V3^ mice demonstrated that the ONL thickness in the medial retina of these mice was not significantly different from that of B6 WT mice (**Fig. 2A, B**). Evaluation of the retinal ultrastructure at the interface between RPE and POS revealed that *Mertk* ^-/- V2^ and *Mertk* ^-/- V3^ mice at 6 months of age had well-preserved RPE microvilli and POS (**Fig. 2C**). Consistent with these histological and ultrastructural findings, retinal function was preserved in 6-month-old *Mertk* ^-/-V2^ and *Mertk* ^-/- V3^ mice as assessed by scotopic electroretinogram recordings (ERGs) (**Fig. 2D-G**). Light-evoked responses in PRs (a-wave) and inner retinal cells (b-wave) in dark-adapted *Mertk* ^-/- V2^ and *Mertk* ^-/- V3^ mice were comparable to those in B6 WT mice, when tested at increasing luminance levels (**Fig. 2D-G**). *Mertk* ^-/- V1^ mice displayed barely any retinal response to light, which is consistent with earlier studies ^4^ and the extensive PR degeneration observed (**Fig. 2D-G**). Thus, the retinal degeneration characteristic of *Mertk* ^-/- V1^ mice is not phenocopied by knocking out *Mertk* in B6 ES cell-derived mice (*Mertk* ^-/- V2^ and *Mertk* ^-/- V3^ mice).

**Figure 2.**
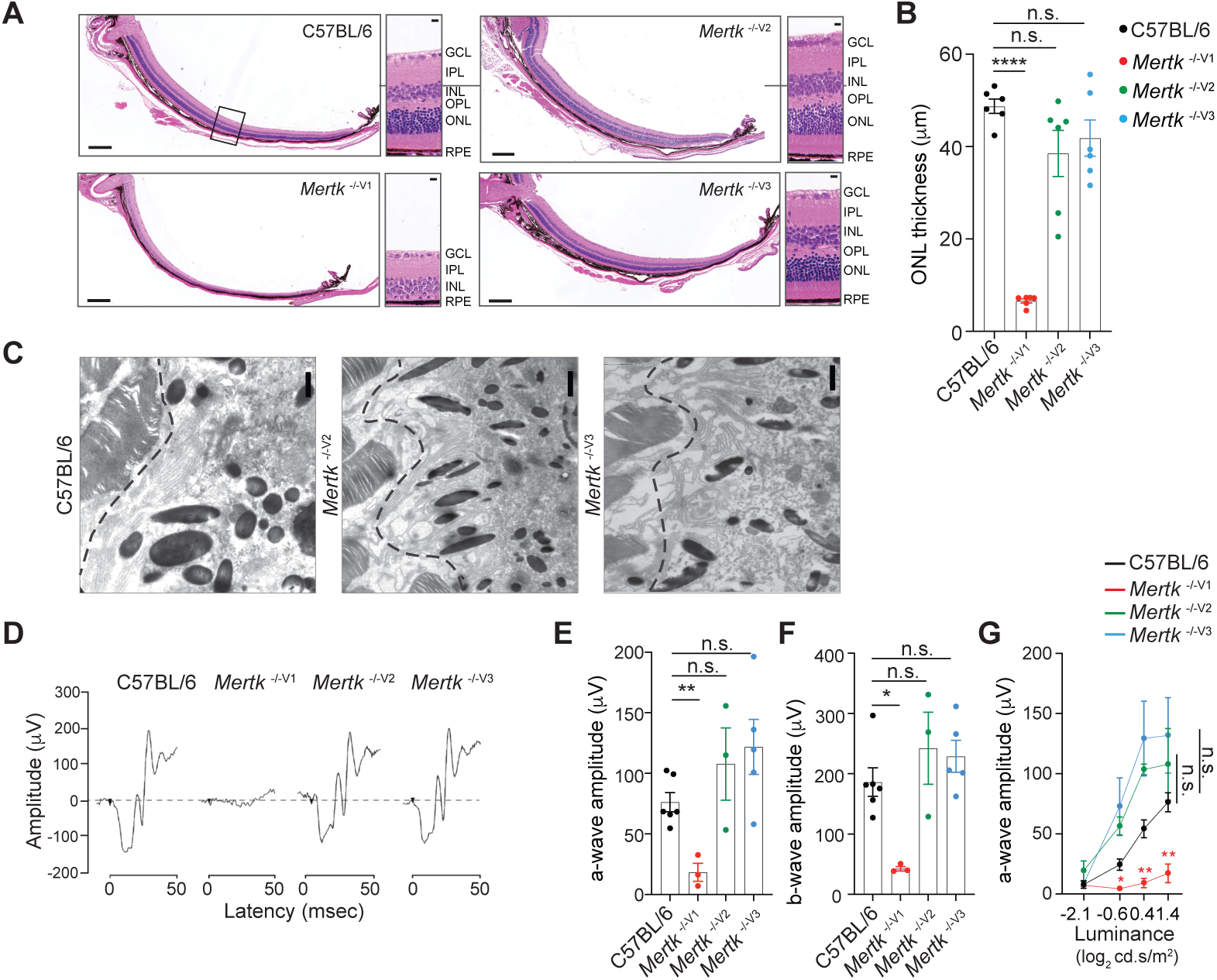
Retinal degeneration of *Mertk* ^-/- V1^ mice is not phenocopied by *Mertk* ^-/- V2^ and *Mertk* ^-/- V3^ mice. Morphological and functional changes in the eye were assessed in 6-month-old C57BL/6, *Mertk* ^-/- V1^, *Mertk* ^-/- V2^ and *Mertk* ^-/- V3^ mice. (**A**) Representative hematoxylin-eosin stained transverse sections of the retina. Boxed section is shown as inset and indicates the area quantified in **B**. Scale bars = 200 μm (left panels) and 10 μm (insets). (**B**) Quantification of outer nuclear layer (ONL) thickness in the area indicated in **A**. Mean ± SEM of 10 measurements/mouse, n=6 mice/genotype. ****p<0.0001, ANOVA-Dunnet’s test. (**C**) Ultrastructure at the POS-RPE interface (dashed line) by transmission electron microscopy. Scale bars = 1 μm. (**D**) Representa- tive scotopic electroretinogram traces are shown at the highest luminance tested. (**E**) Quantification of a-wave amplitude at highest luminance tested (25 cd.s/m^2^), n=3-6 mice/ genotype. **p<0.01, ANOVA-Dunnet’s test. (**F**) Quantification of b-wave amplitude at the highest luminance tested (25 cd.s/m^2^). *p<0.05, ANOVA-Dun- net’s test. (**G**) a-wave amplitude at increasing luminances. *p<0.05, **p<0.01, 2-way ANOVA.

### *Mertk* targeting in 129P2 ES cells results in a robust anti-tumor response against immune checkpoint inhibitor (ICI)-refractory YUMM1.7 melanoma and GL261 brain tumor, but this effect is not phenocopied in *Mertk* ^-/- V2^ or *Mertk* ^-/- V3^ mice

An important, newly discovered role of MERTK is as an innate immune checkpoint in cancer ^10,12,13,21-27^. *Mertk* ^-/- V1^ mice were used to demonstrate improved anti-tumor immune response against MMTV-PyVmT, B16:F10 and MC38 tumor models ^10^. Based on the use of *Mertk* ^-/- V1^ mice, it was also surmised that inhibition of macrophage efferocytosis due to MERTK loss-of-function results in decreased tumor growth and increased tumor-free survival in E0771 murine breast cancer model ^12^. Similarly, *Mertk* ^-/- V1^ mice were used in a study by Lindsay *et al*. to demonstrate improved anti-tumor T cell motility in a B78ChOva tumor model ^13^. We compared the anti-tumor response of *Mertk* ^-/- V1^, *Mertk* ^-/- V2^ and *Mertk* ^-/- V3^ mice in a model of ICI-refractory melanoma (YUMM1.7), as well as in an orthotopic brain tumor mouse model (GL261). YUMM1.7 tumor cells were implanted subcutaneously in *Mertk* ^-/- V1^, *Mertk* ^-/- V2^ and *Mertk* ^-/- V3^ mice or in B6 WT (**Fig. 3A**). Tumor-free survival (TFS), overall survival (OS) and rate of tumor growth were monitored (**Fig. 3A, B**). Consistent with previous reports of MERTK blockade enhancing anti-tumor response ^10,12,13,21-27^, 100% of *Mertk* ^-/- V1^ mice remained tumor free, compared to 0% of B6 WT control mice (**Fig. 3B**). Surprisingly, neither of the *Mertk* knockout mice generated using B6 ES cells, *Mertk* ^-/- V2^ and *Mertk* ^*-/-* V3^, phenocopied the tumor resistance observed in *Mertk* ^-/- V1^ mice. 100% of *Mertk* ^-/- V2^ and *Mertk* ^*-/-* V3^ mice succumbed to tumor growth and at a rate comparable to their B6 WT counterparts (**Fig. 3B**). To investigate these findings in an independent tumor model, luciferase expressing GL261 brain tumor cells were orthotopically injected in B6 and *Mertk* ^-/- V1^ mice. Bioluminescence imaging was performed to determine tumor volume and mice were monitored for OS (**Fig. S2**). No differences were observed in the rate of GL261 growth or time to end-point between B6 WT or *Mertk* ^-/- V1^ mice (**Fig S2B, C**). We next attempted to treat tumor-bearing mice with a dendritic cell vaccine (DC-Vax). Tumor cell lysates obtained by freeze-thawing GL261 were fed to B6 WT bone marrow-derived DCs. Following co-incubation, DCs were activated with LPS treatment and intraperitoneally injected into tumor-bearing B6 WT or *Mertk* ^-/- V1^ mice on day 14 and day 21 after tumor implantation (**Fig. 3C**). Of note, tumor sizes were comparable between B6 WT or *Mertk* ^-/- V1^ mice prior to receiving the DC-vax (**Fig. 3E**, top panel). Remarkably, the administration of a DC-Vax in *Mertk* ^-/- V1^ mice resulted in a significant reduction in tumor size, whereas DC-Vax-treated B6 WT mice succumbed to their tumor burden (**Fig. 3D, E**). Similar to the studies on YUMM1.7 melanoma cells, when GL261 cells were implanted in *Mertk* ^-/- V2^ and *Mertk* ^-/- V3^ mice, and these mice were subsequently treated with DC-Vax, 100% of these mice failed to display the anti-GL261 response of *Mertk* ^-/- V1^ mice (**Fig. 3D, E**). Taken together, our experiments revealed an astounding anti-tumor resistance of *Mertk* ^-/- V1^ mice against both YUMM1.7 and GL261. However, this phenotype of *Mertk* ^- /- V1^ mice was again not phenocopied by *Mertk* ^-/- V2^ and *Mertk* ^-/- V3^ mice.

**Figure 3.**
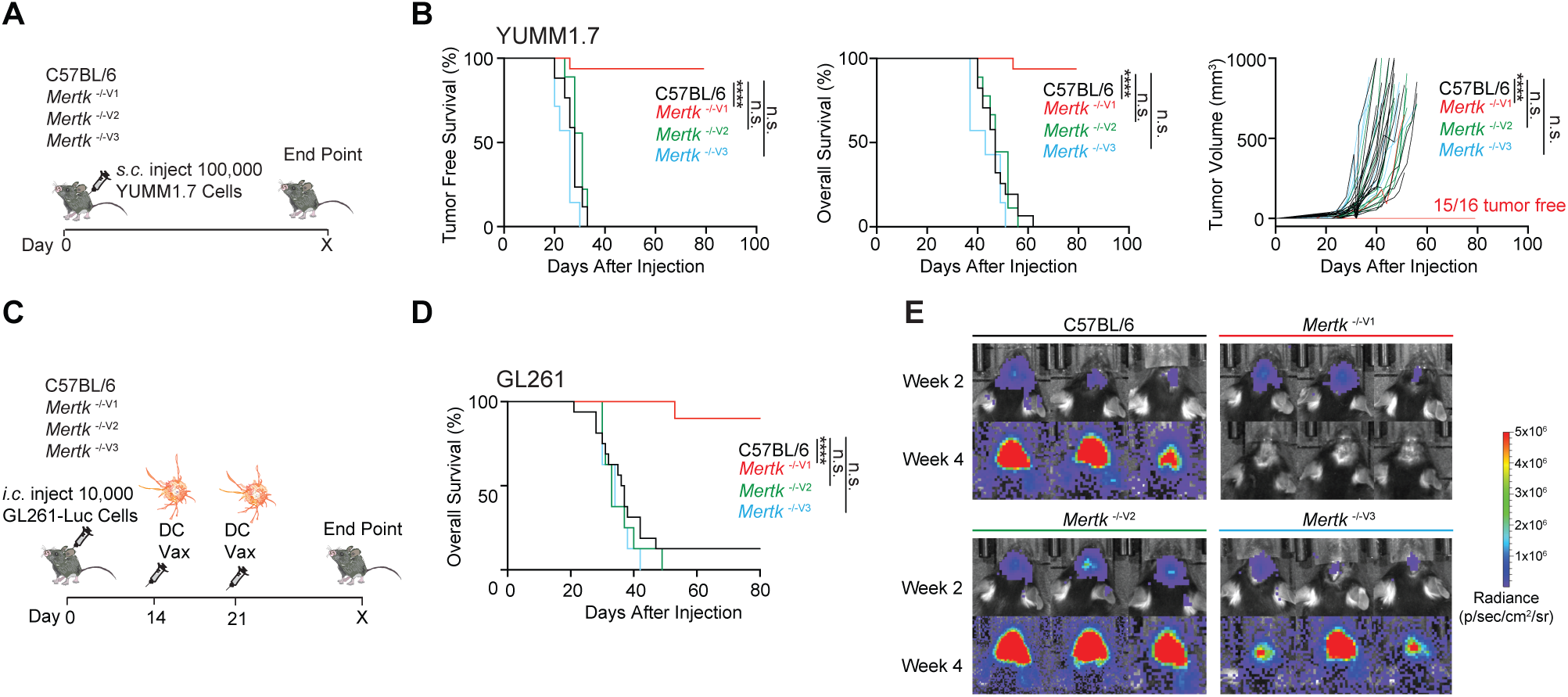
Anti-tumor response of *Mertk* ^-/- V1^ mice is not phenocopied by *Mertk* ^-/- V2^ and *Mertk* ^-/- V3^ mice. (**A**) Schematic showing subcutaneous injection of 100,000 *Braf* ^V600E^ *Pten* ^-/-^ YUMM1.7 mouse melanoma cells into mice of different genotypes. (**B**) Tumor free survival (TFS), overall survival (OS) and tumor volume in C57BL/6 (n=17), *Mertk* ^-/- V1^ (n=16), *Mertk* ^-/- V2^ (n=9) and *Mertk* ^-/- V3^ mice (n=7) mice implanted with YUMM1.7 cells. ****p<0.0001, Log-rank Mantel-Cox test or two-way ANOVA, Dunnet’s mutliple comparison test. (**C**) Schematic showing intracranial injection of 10,000 GL261-Luc glioma cells and intraperitoneal dendritic cell vaccination of mice at 14 and 21 days post-tumor implantation. (**D**) OS in C57BL/6 (n=16), *Mertk* ^-/- V1^ (n=10), *Mertk* ^-/- V2^ (n=8) and *Mertk* ^-/- V3^ (n=8) mice implanted with GL261 tumors. ****p<0.0001, Log-rank Mantel-Cox test. (**E**) Representative IVIS images of intracranial tumors at D14 and D28 post-implantation.

### Gene expression differences revealed by genome-wide transcriptional analyses in the presence of 129P2 *versus* B6 alleles on chromosome 2

A segment of chromosome 2 in the *Mertk* ^-/- V1^ mice was previously reported to be derived from the 129P2 background ^19^. Therefore, we performed short tandem repeat (STR) analysis in the chromosome 2 region surrounding *Mertk* locus in B6 WT, *Mertk* ^-/- V1^, *Mertk* ^-/- V2^ and *Mertk* ^-/- V3^ mice. Genomic DNA isolated from each of these mouse lines was subjected to PCR amplification of 24 microsatellite sites across chromosome 2 **(Fig. S3)**. We found that *Mertk* ^-/- V1^ mice harbored a 15.08 cM region between D2Mit206 and D2Mit168 that is of 129P2 origin (**Fig. 4A**). As expected, chromosome 2 was entirely B6-derived in both *Mertk* ^-/- V2^ and *Mertk* ^-/- V3^ mice **(Fig. 4A)**.

**Figure 4.**
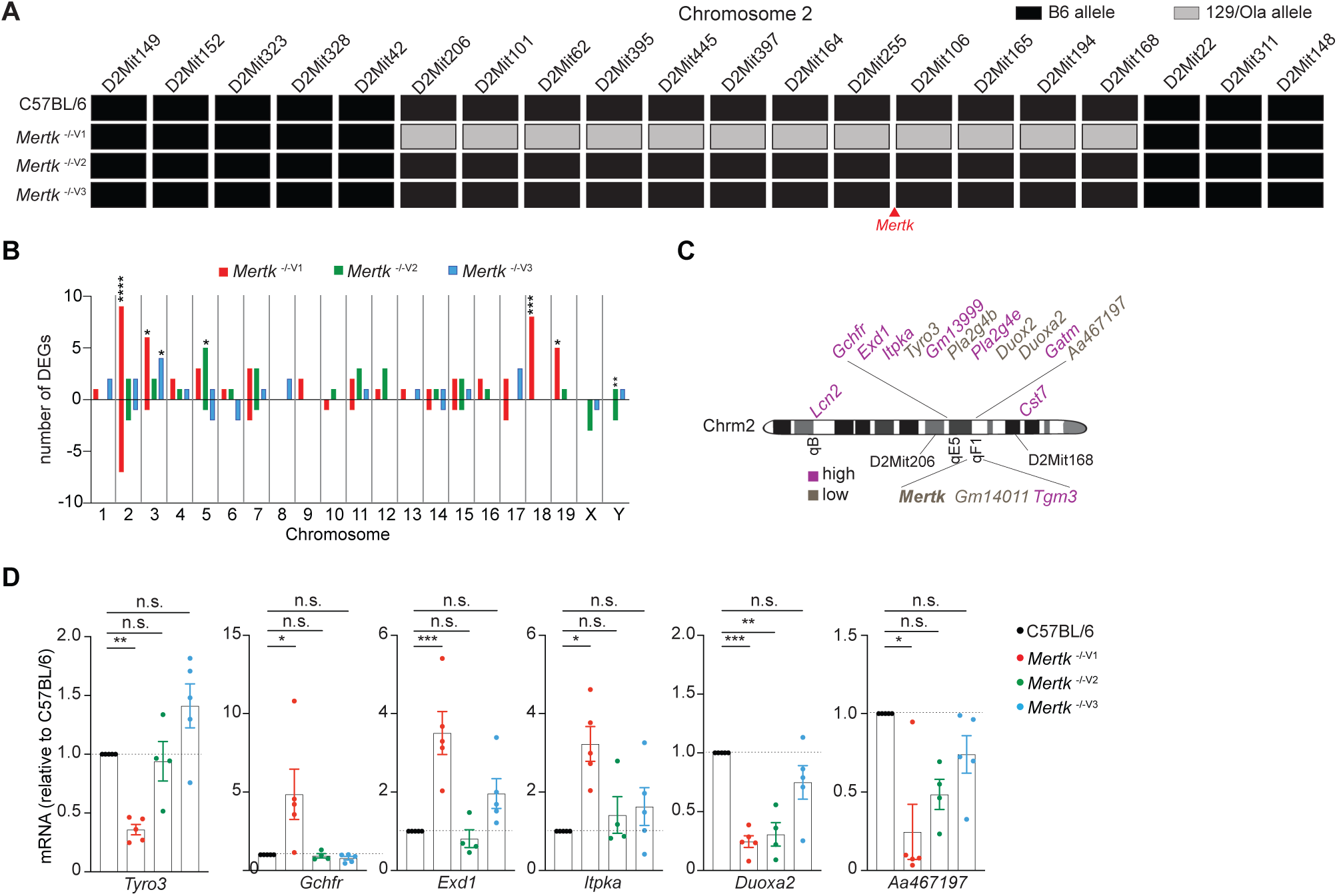
Significant gene expression differences in RPE of *Mertk* ^-/- V1^ versus *Mertk* ^-/- V2^ and *Mertk* ^-/- V3^ mice. (**A**) Haplotype map comparing 20 microsatellite markers across chromosome 2 in C57BL/6, *Mertk* ^-/- V1^, *Mertk* ^-/- V2^ and *Mertk* ^-/- V3^ mice. Black rectangles indicate homozygosity for C57BL/6 (B6) alleles and gray rectangles indicate homozygosity for 129P2/OLAHsd (129/Ola) alleles. (**B**) Distribution of DEGs in RPE across chromosomes. *p<0.05, **p<0.01, ***p<0.001, ****p<0.0001, hypergeometric test. (**C**) Schematic showing genes neighboring *Mertk* that are significantly upregulated or downregulated in the RPE of *Mertk* ^-/- V1^ mice. (**D**) qPCR quantification of indicated chro- mosome 2 genes in C57BL/6, *Mertk* ^-/- V1^, *Mertk* ^-/- V2^ and *Mertk* ^-/- V3^ RPEs. *p<0.05, **p<0.01, ***p<0.001, one-way ANOVA-Dunnet’s test.

To more broadly understand the genome-wide transcriptional differences between *Mertk* ^-/- V1^, *Mertk* ^-/- V2^ and *Mertk* ^-/- V3^ mice accounting from the chromosomal differences, we performed RNA sequencing (RNAseq) experiments on RPE from these three mouse lines at postnatal day (P) 25. We identified a number of genes that were differentially expressed in *Mertk* ^-/- V1^, *Mertk* ^-/- V2^ and *Mertk* ^-/- V3^ RPE in *cis* and in *trans* compared to B6 WT RPE (**Fig. 4B**). Notable amongst the changes in *cis* in *Mertk* ^-/- V1^ RPE was *Tyro3* (**Fig. 4C**). Significant changes in the transcripts of *Tyro3* and other *Mertk*-neighboring genes, in *Mertk* ^-/- V1^, but not *Mertk* ^-/- V2^ and *Mertk* ^-/- V3^ RPE, relative to B6 WT RPE were confirmed by qPCR (**Fig. 4D**).

### *Tyro3* is epistatic with *Mertk* for the retinal degeneration trait

It was previously reported that TYRO3 levels were significantly lower in *Tyro3*^129/129^ RPE compared to *Tyro3*^B6/B6^ or *Tyro3*^129/B6^ RPE ^19^. Consistent with this report, we detected significantly lower levels of TYRO3 in *Mertk* ^-/- V1^ RPE (**Fig. 5A, B**). By contrast, RPE cells from *Mertk* ^-/- V2^ and *Mertk* ^-/- V3^ mice had levels of TYRO3 that were comparable to B6 WT mice (**Fig. 5A, B**). Since it was concluded that even the hypomorphic expression of *Tyro3*^B6/129^ can suppress the phenotypes of *Mertk* loss-of-function ^19^, we tested whether the simultaneous genetic ablation of *Mertk* and *Tyro3* can phenocopy *Mertk* ^-/- V1^ mice. We engineered *Mertk* ^-/- V2^ *Tyro3* ^-/- V2^ mice by targeting *Tyro3* in *Mertk* ^-/- V2^ mice using CRISPR/CAS9 (**Fig. 5C, Fig. S1D**). Immunoblotting experiments confirmed that TYRO3 was undetectable in the RPE of *Mertk* ^-/- V2^ *Tyro3* ^-/- V2^ mice, when compared to TYRO3 expression in B6 WT RPE (**Fig. 5D, E**). Indeed, these mice recapitulated the severe retinal degeneration observed in *Mertk* ^-/- V1^ underscoring the function of *Tyro3* ^B6^ as a suppressor allele in retinal degeneration induced by targeting *Mertk*. When eye sections from 6-months old *Mertk* ^-/- V2^ *Tyro3* ^-/- V2^, *Mertk* ^-/- V1^ and B6 WT controls were stained with hematoxylin and eosin, we found that ONL thickness in *Mertk* ^-/- V2^ *Tyro3* ^-/- V2^ mice was significantly reduced compared to B6 WT controls, commensurate with *Mertk* ^-/- V1^ mice (**Fig. 5F, G**). Similar to *Mertk* ^-/- V1^ mice, *Mertk* ^-/- V2^ *Tyro3* ^-/- V2^ mice had only ∼1 row of nuclei in the ONL across the entire dorsal-ventral axis of the retina (**Fig. 5F, G**). Consequently, *Mertk* ^-/- V2^ *Tyro3* ^-/- V2^ mice did not display light-evoked responses in scotopic ERGs at any luminance tested (**Fig. 5H, I**). These results show that *Mertk* ^-/- V2^ *Tyro3* ^-/- V2^ mice phenocopy the widespread morphological and functional retinal deficits characteristic of *Mertk* ^-/- V1^ mice.

**Figure 5.**
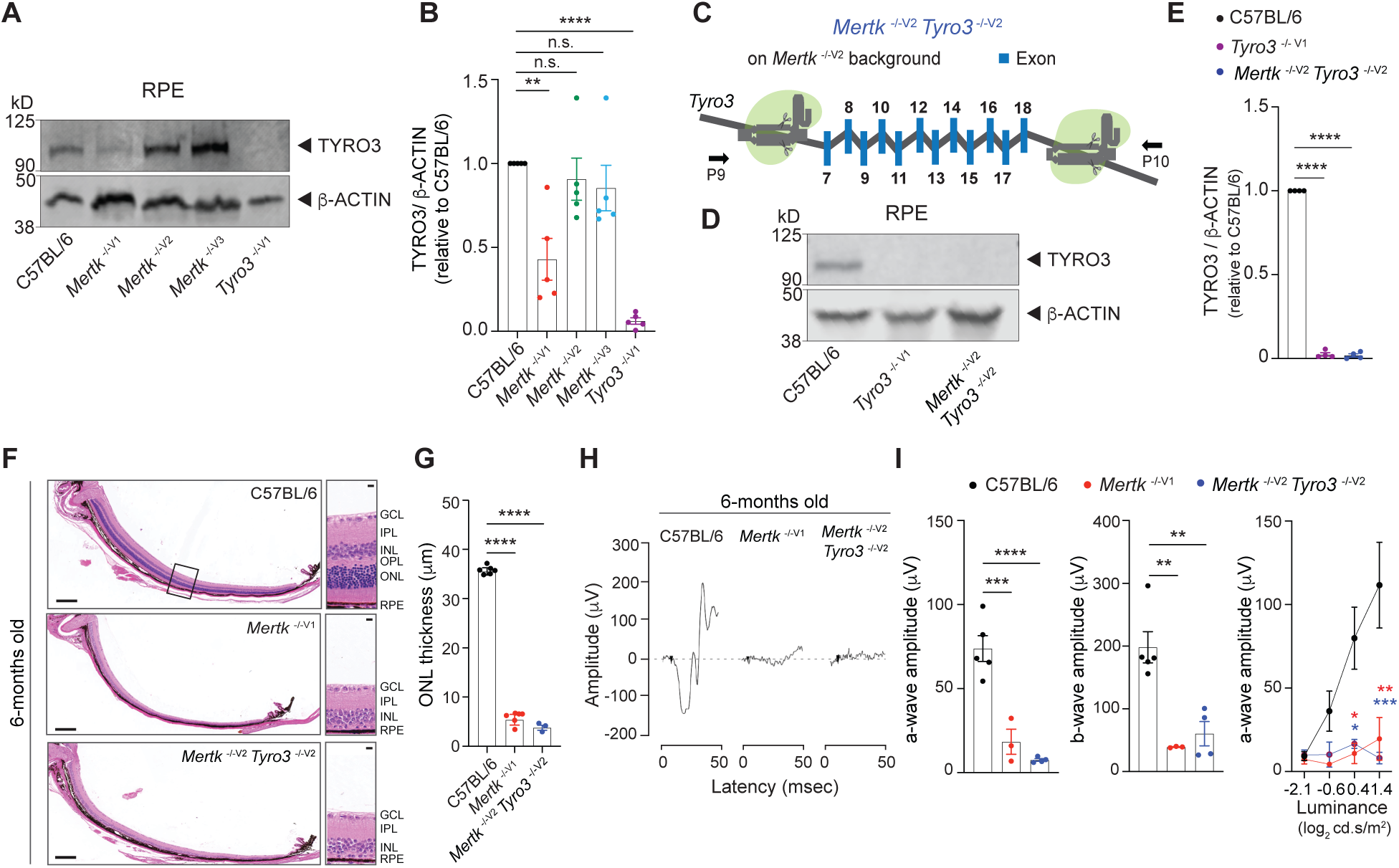
*Tyro3* is epistatic with *Mertk* for the retinal degeneration trait. (**A, B**) Representative and independent measurements of TYRO3 amounts in RPE. Western blot (WB) from C57BL/6, *Mertk* ^-/- V1^, *Mertk* ^-/- V2^, *Mertk* ^-/- V3^ and Tyro3 ^-/- V1^ mice RPE (mean ± SEM of n=5 mice/ genotype). *p<0.05, **p<0.01, ***p<0.001 and ****p<0.0001, one-way ANOVA-Dunnet’s test. (**C**) Schematic showing targeting of *Tyro3* exons 7-18 with CRISPR/Cas9 in *Mertk* ^-/- V2^ ES cells to generate the *Mertk* ^-/- V2^ *Tyro3* ^-/- V2^ mouse line. Image not drawn to scale. (**D, E**) Representative and independent measurements of TYRO3 amounts in RPE. WB from C57BL/6, *Mertk* ^-/- V2^ *Tyro3* ^-/- V2^ and *Tyro3* ^-/- V1^ mice RPE (mean ± SEM of n=3 mice/ genotype). ****p<0.0001, one-way ANO- VA-Dunnet’s test. (**F**) Representative hematoxylin-eosin stained transverse sections of the retina. Boxed section is shown as inset and indicates the areas quantified in (**G**). Scale bars = 200 μm (left panels) and 10 μm (insets). (**G**) Quantification of outer nuclear layer (ONL) thickness in the area indicated in **F**. Mean ± SEM of 10 measurements/mouse, n=6 mice/genotype. ****p<0.0001, one-way ANOVA-Dunnet’s test. (**H**) Representative scotopic electroretinogram traces are shown at the highest luminance tested. (**I**) Quantification of a-wave amplitude and b-wave at highest luminance tested, n=3-6 mice/genotype. **p<0.01, ***p<0.001, ****p<0.0001, ANOVA-Dunnet’s test. a-wave amplitude at increasing luminances. *p<0.05, **p<0.01, ***p<0.001, two-way ANOVA. Morphological and functional changes in the eye were assessed in 6-month-old C57BL/6, *Mertk* ^-/-V1^ and *Mertk* ^-/- V2^ *Tyro3* ^-/- V2^ mice.

### Neither deficient efferocytosis in macrophages nor loss of *Tyro3* ^B6/B6^ can universally account for the anti-tumor immunity in *Mertk* ^-/- V1^ mice

The prevailing dogma is that the absence of MERTK in tumor-associated macrophages prevents the phagocytosis and disposal of dead tumor cells, thereby increasing the availability of tumor-antigen to DCs and triggering improved anti-tumor T cell immunity ^12^. Alternatively, deficient MERTK-dependent phagocytosis by tumor-associated macrophages leads to secondary necrosis of tumor cells and increased pro-inflammatory microenvironment more conducive to anti-tumor immunity ^11^. Of note, these scenarios are not mutually exclusive and might in fact cooperate. Nevertheless, deficient phagocytosis of tumor cells by macrophages in the absence of *Mertk* is described as the *sine qua non* of the remarkable anti-tumor immunity observed when MERTK function is either genetically ablated or pharmacologically inhibited ^11,12,23^. We performed similar RNAseq analyses in BMDMs isolated from *Mertk* ^-/- V1^, *Mertk* ^-/- V2^ and *Mertk* ^-/- V3^ and B6 WT mice. Our experiments identified a number of changes in *Mertk* ^-/- V1^, *Mertk* ^-/- V2^ and *Mertk* ^-/- V3^ BMDMs, compared to B6 WT controls. Furthermore, *Mertk* ^-/- V1^ mice had a significant number of upregulated and downregulated genes that were not recapitulated in *Mertk* ^- /- V2^ and *Mertk* ^-/- V3^ mice (**Fig. 6A-C**). Albeit that the lack of anti-tumor resistance in *Mertk* ^-/- V2^ and *Mertk* ^-/- V3^ mice already pointed to a MERTK agnostic basis for tumor clearance, we hypothesized that some of the coincidental changes in BMDMs in these mice might compensate for the loss of MERTK by providing redundancy in phagocytosis. Thus, we expected no reduction in efferocytosis in BMDMs derived from *Mertk* ^-/- V2^ and *Mertk* ^-/- V3^ mice, unlike BMDMs from *Mertk* ^-/- V1^ mice. We generated CD11b^+^ F4/80^+^ BMDMs from B6 WT, *Mertk* ^-/- V1^, *Mertk*^-/- V2^ and *Mertk* ^-/- V3^ mice and tested them in an *ex vivo* phagocytosis assay. Flow cytometry-based analysis of BMDMs derived from *Mertk* ^-/- V2^ and *Mertk* ^-/- V3^ mice confirmed MERTK ablation in these cells, compared to B6 controls **(Fig. S4A, B)**. BMDMs were cultured with pHrodo-labeled apoptotic thymocytes in the presence of serum and their uptake was assayed by flow cytometry after 1 hr. As expected, *Mertk* ^-/- V1^ BMDMs were ∼50% less phagocytic than B6 WT BMDMs. BMDMs from *Mertk* ^-/- V2^ and *Mertk* ^-/- V3^ mice were similarly less phagocytic than those from B6 WT mice **(Fig. 6D, E)**. These experiments demonstrate that BMDMs derived from *Mertk* knock out mice of 129P2 or B6 origin were equally deficient in the efferocytosis of apoptotic thymocytes.

**Figure 6.**
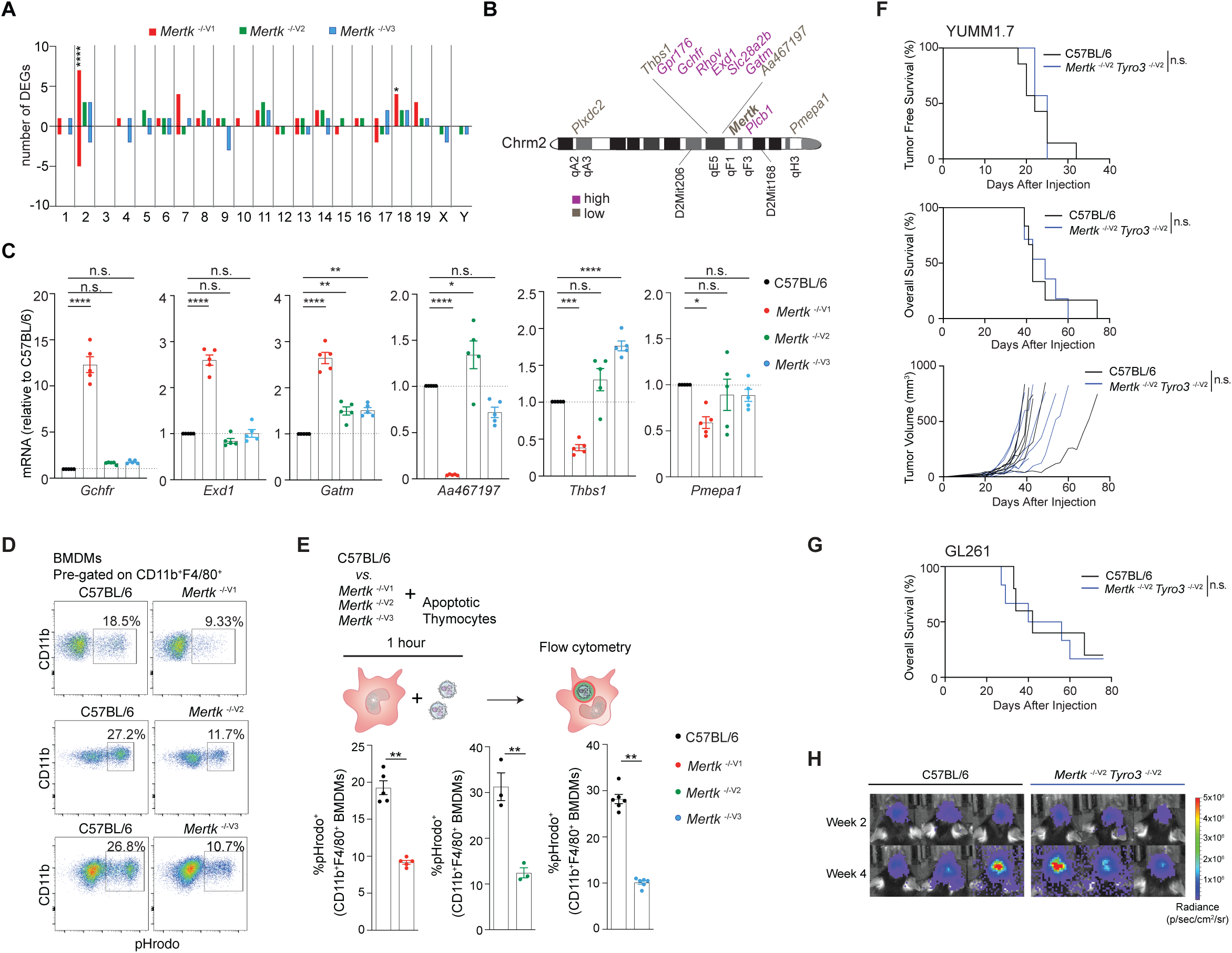
Anti-tumor response of *Mertk* ^-/- V1^ mice is neither the result of deficient efferocytosis by macrophages nor hypomorphic TYRO3. **(A)** Distribution of differentially expressed genes (DEGs) in bone-marrow-derived-macrophages (BMDMs) across chromosomes. *p<0.05, ****p<0.0001, hypergeometric test. (**B**) Schematic indicating chromosome 2 genes that are upregulated or downregulated in *Mertk* ^-/-V1^ BMDMs. (**C**) qPCR quantification of the indicated chromosome 2 genes in C57BL/6J, *Mertk* ^-/- V1^, *Mertk* ^-/- V2^ and *Mertk* ^-/- V3^ BMDMs. *p<0.05, **p<0.01, ***p<0.001, ****p<0.0001, one-way ANOVA-Dunnet’s test. (**D**) BMDMs from C57BL/6, *Mertk* ^-/- V1^, *Mertk* ^-/- V2^ and *Mertk* ^-/- V3^ mice were co-cultured with apoptotic thymocytes for 1 hour. Representative plots show uptake of pHrodo labeled apoptotic thymocytes by CD11b^+^F4/80^+^ BMDMs by flow cytometry. (**E**) Quantification of **D** (n=3-6 mice/group, **p<0.01, Mann-Whitney or student t-test). (**F**) Tumor-free survival, overall survival (OS) and tumor volume in C57BL/6 (n=7), *Mertk* ^-/-V1^ (n=7) and *Mertk* ^-/- V2^ *Tyro3* ^-/- V2^ (n=7) implanted with YUMM1.7 tumors. n.s., Log-rank Mantel-Cox test. (**G**) OS in C57BL/6 (n=5) and *Mertk* ^-/- V2^ *Tyro3* ^-/- V2^ (n=6) mice implanted with GL261 tumors. n.s., Log-rank Mantel-Cox test. (**H**) Representative IVIS images of intracranial tumors in C57BL/6 and *Mertk* ^-/- V2^ *Tyro3* ^-/- V2^ mice at D14 and D28 post-implantation.

It is important to note that TYRO3 can not only compensate for MERTK in phagocytosis ^19^, it is also a negative regulator of the immune response ^28^. However, *Tyro3* was not a differentially expressed gene (DEG) in BMDMs because it is not expressed in these cells (**Fig. 6A-C**). To entirely rule out *Tyro3* ^B6^ function as a suppressor in abolishing the anti-tumor effects of *Mertk* ablation in *Mertk* ^-/- V2^ and *Mertk* ^-/- V3^ mice, we implanted YUMM1.7 and GL261 tumors in *Mertk* ^-/- V2^ *Tyro3* ^-/- V2^ mice to dispense off the *Tyro3* redundancy and potentially reveal the anti-tumor resistance. However, we observed that *Mertk* ^-/- V2^ *Tyro3* ^-/- V2^ mice entirely failed to phenocopy the anti-tumor resistance observed in *Mertk* ^-/- V1^ mice (**Fig. 6F-H**). Specifically, 100% of *Mertk* ^-/- V2^ *Tyro3* ^-/- V2^ mice failed to show improved survival in comparison to B6 WT mice when implanted with YUMM1.7 (**Fig. 6F**). No differences were observed in TFS, OS or in tumor growth (**Fig. 6F**). Similarly, in the GL261 model, no differences were observed in OS or in tumor volume following the DC vaccination strategy that conferred significant anti-tumor resistance to *Mertk* ^-/- V1^ mice (**Fig. 6G, H**).

### Tissue-specific discordance in gene expression between *Mertk* ^-/-^ mouse lines carrying 129P2 *versus* B6 alleles

How are *Mertk* ^-/- V1^ mouse traits modified? RNAseq demonstrated that gene expression changes in *Mertk* ^-/- V1^ mice are not restricted to changes in *cis* on chromosome 2, but expanded in *trans* to other chromosomes. A number of genes displaying differential expression in *Mertk* ^-/- V1^ *versus Mertk* ^-/- V2^ and *Mertk* ^-/- V3^ in BMDMs mapped between D2Mit206 [55.94 cM] and D2Mit168 [71.02 cM] (in *cis*) on chromosome 2 (**Fig. 6B**). We were able to confirm that genes mapping to this region, including *Gchfr, Exd1* and *Gatm* were increased by ∼2.5 to 12-folds in *Mertk* ^-/- V1^ but not *Mertk* ^-/- V2^ and *Mertk* ^-/- V3^ BMDMs, compared to B6 WT BMDMs, by qPCR (**Fig. 6C**). Additionally, transcripts such as *Aa467197, Thbs1* and *Pmepa1* were found to be ∼40-80% decreased in *Mertk* ^-/- V1^ but not *Mertk* ^-/- V2^ and *Mertk* ^-/- V3^ BMDMs, compared to B6 WT BMDMs (**Fig. 6C**). Interestingly, genes beyond this chromosomal segment (*i.e*. in *trans*) were also differentially expressed between B6 WT, *Mertk* ^-/- V1^, *Mertk* ^-/- V2^ and *Mertk* ^-/- V3^ BMDMs (**Fig. 6A**). The changes in the expression of genes *in trans* common to *Mertk* ^-/- V1^, *Mertk* ^-/- V2^ and *Mertk* ^-/- V3^ BMDMs by comparison to B6 WT are likely direct downstream consequences of *Mertk* ablation. Nevertheless, there were also distinct changes in *trans* in *Mertk* ^-/- V1^ BMDMs not reproduced in *Mertk* ^- /- V2^ and *Mertk* ^-/- V3^ BMDMs (**Fig. 6A**). Such changes are likely independent of *Mertk*.

Importantly, many of the DEGs in *Mertk* ^-/- V1^ *versus Mertk* ^-/- V2^ and *Mertk* ^-/- V3^ in *cis* as well as in *trans* were distinct between BMDMs and RPE (**Fig. 6A-C and Fig. 4B-D**). For instance, DEGs corresponding to chromosome 2 such as *Iptka, Gm13999, Pla2g4e, Duox2, Duoxa2, Gpr176, Rhov* or *Slc28a2b* were unique to between BMDMs or RPE. Similarly, in chromosome 18 and 19, there were a number of DEGs unique to BMDMs or RPE (**Fig. 6A and Fig. 4B**). Therefore, it is likely that there might be tissue-specific modifiers operating both in *cis* and *trans*. Thus, complex traits in *Mertk* ^-/- V1^ mice, such as anti-tumor resistance, are likely to result from a combination of cell type-specific changes.

## DISCUSSION

Generation of the *Mertk* ^-/- V1^ mice ^2^ was instrumental for functional studies on this RTK revealing its critical role in phagocytosis and negative regulation of inflammation ^2,3^. This mouse model, heretofore, has remained the workhorse for investigation of the biological roles of *Mertk*. There is an increasing contemporary interest in the role of this RTK in what are likely to be complex traits, such as anti-tumor response and neurodegeneration ^5-7,10-15^. Loss of/reduced phagocytosis by macrophages and other phagocytes, when *Mertk* is genetically ablated, is often surmised causal to the manifested phenotype ^7,11,12,23^. The functional role of MERTK in phagocytosis remains beyond aspersions and we, in fact, validated the dependency of phagocytosis by macrophages on MERTK using two independently generated mouse knockout lines. In mice wherein MERTK signaling was disabled by targeting the sequence coding for the kinase domain of MERTK (*Mertk*^K614M/K614M^ mice), peritoneal macrophages failed to phagocytose apoptotic thymocytes at rates similar to that observed in *Mertk* ^-/- V1^ mice. Importantly, *Mertk*^K614M/K614M^ mice were generated in B6 background, further indicating that the phagocytic function of macrophages depended on MERTK and is lost in its absence, independent of the 129P2 or B6 ES cell background *Mertk* was targeted in ^29^. Neither do we consider all *Mertk* loss-of-function traits to be ambiguous. Using *Mertk* ^-/- V1^ mice, DeBerge *et al*. demonstrated that post-reperfusion MERTK-dependent phagocytosis is cardioprotective in a mouse model of myocardial ischemia reperfusion injury ^30^. Notably, conditional ablation of *Mertk* in myeloid cells of B6 background confirmed a cardioprotective role for this RTK ^30^. Yet, many phenotypes of *Mertk* ^-/- V1^ mice are not invariably and inviolably linked to loss of phagocytosis in the absence of MERTK. This includes, at the very least, retinal degeneration and anti-tumor immunity.

Here we identify the existence of independent modifiers of *Mertk* loss-of-function traits, including *Tyro3*. The original *Mertk* ^-/-^ mouse line, even after >10 generations of backcrossing to B6 mice in our laboratory, retained a ∼15 cM chromosome 2 segment with 129P2 alleles that are genetically linked to the targeted *Mertk* locus and failed to recombine and segregate from it. A previous report by Vollrath *et al*. had also reported the presence of a ∼40 cM chromosome 2 segment with 129P2 alleles in *Mertk* ^-/- V1^ mice ^19^. Vollrath *et al*. mapped a modifier of the retinal degeneration phenotype to a 2 cM region containing 53 ORFs by genetic linkage mapping ^19^. Segregation of the *Mertk*-linked 129 region with a B6 region that contained *Tyro3* restored retinal homeostasis ^19^. Since a genetic strategy was used in this previous report, in theory, potential contribution of other genes within the co-segregating region with suppressor function cannot be ruled out. Given that *Tyro3* is the only *Mertk* paralog within this region, such an exception is unlikely. Nonetheless, in a complementary approach, we directly targeted *Tyro3* in B6 *Mertk* ^-/- V2^ mice-derived ES cells to unambiguously demonstrate that the simultaneous ablation of *Mertk* and *Tyro3* in B6 mice is necessary and sufficient for retinal degeneration. One of the potential concerns for using MERTK inhibitors in the clinic is the on target adverse event of rapid retinal degeneration. Vollrath *et al*. ^19^, Maddox *et al*.^16^ and our results collectively indicate that sparing TYRO3 activity can greatly ameliorate this concern while targeting the oncogenic activity of MERTK.

There is an emerging concept of MERTK as an innate immune checkpoint and that targeting this RTK might enhance immunotherapy ^22,31-34^. Remarkable anti-tumor resistance was observed in a number of standard mouse tumor models such as MMTV-PyVmT, B16:F10, MC38, E0771, B78ChOva in *Mertk* ^-/- V1^ mice ^10,12,13^. Our initial results were entirely consistent with and extended such findings in that we were able to observe an astonishing anti-tumor resistance in hard-to-treat mouse models such as the anti-CTLA-4- and anti-PD-1-refractory YUMM1.7 mouse melanoma. *Mertk* ^-/- V1^ mice could even be rendered resistant to the orthotopic GL261 with post-tumor treatment with DCvax. Although our use of additional tumor models further validated host anti-tumor resistance in the *Mertk* ^-/- V1^ mice paradigm, we surprisingly also discovered that *Mertk* ablation is not sufficient for resistance against YUMM1.7 and GL261. Nevertheless, it is important to acknowledge that the work by Davra *et al*. and Lindsay *et al*. not only employed the *Mertk* ^-/- V1^ mouse, but also utilized alternative approaches such as antibodies or small molecule inhibitors to pharmacologically disable MERTK ^12,13^. The role of *Mertk* as an oncogene confounds interpretations of ostensible immune function when using pharmacological agents. It is possible that some of the reported effects are a consequence of inhibiting the oncogenic role of TAM RTKs. Since we did not test identical tumor models as the previous reports, it remains entirely possible that anti-tumor immunity in the models reported before, serendipitously, are dependent exclusively on the loss of MERTK. However, the universality of improving anti-tumor immunity irrespective of tumor models employed through targeting MERTK has to be questioned. Phenotypic differences not accounted for fully by the target gene are not entirely an uncommon occurrence in tumor resistance. For example, the frequency of intestinal adenomas in *Apc*^min^ mice is dramatically modified by the genetic background that the mutation is introduced in. *Apc*^min^ mice with a B6 background were reported to develop 28.5±7.9 tumors and die within 4 months of birth ^35^. When these *Apc*^min^ mice were crossed to an AKR mouse, the mixed B6/AKR background had 5.8±4.3 tumors and lived till they were euthanized at 300 days ^35^. Crossing the B6/AKR F1 mice with AKR further reduced tumor loads to 1.75±1.7 tumors ^35^. The modifier allele was identified as *Mom1* mapping to chromosome 4 ^35^. We postulate that the anti-tumor resistance of *Mertk* ^-/- V1^ mouse line also involves modifier alleles. The 129P2 segment in *Mertk* ^-/- V1^ mice not only accounts for a number of gene expression differences within the *cis* non-B6 flanking region, but also results in differential gene expression in a number of *trans* loci. It is possible that the anti-tumor resistance ascribed solely to *Mertk* might actually involve one or more modifier activities encoded within *cis* or *trans* loci unique to the *Mertk* ^-/- V1^ mice that are absent in B6 ES cell-derived *Mertk* ^-/-^ mouse lines. Furthermore, since gene expression differences between *Mertk* ^-/- V1^ and *Mertk* ^-/- V2^ or *Mertk* ^-/- V3^ mice are cell type-specific, it is possible that tissue-specific expression quantitative trait loci (eQTLs) may function in combination as modifiers of complex traits, such as anti-tumor immune response. This makes identification of the crucial modifiers more challenging. Assuming said modifiers are acting in *cis*, genomic CRISPR/CAS9 screens for loss of YUMM1.7 and GL261 resistance in a series of *Mertk* ^-/- V1^-derived mouse lines ablated for genes within the ∼15 cM region of chromosome 2 may reveal the modifiers. Alternatively, if the 129P2 allele is dominant, BAC transgenics using *Mertk* ^-/- V2^ ES cells to express 129P2 variant of genes for the rescue of the anti-tumor response might also be an appropriate strategy.

Several gene expression changes were observed in *Mertk* ^-/- V1^ *versus Mertk* ^-/- V2^ and *Mertk* ^-/- V3^ RPE. We performed RNAseq at P25 because the retinal degeneration characteristic of our *Mertk* ^-/- V1^ mouse colony is not yet evident at this age and therefore, the changes are unlikely to be a consequence of retinal degeneration. Consistent with this notion, the changes were concentrated on chromosome 2 and therefore linked to the 129P2 origin of the alleles. The biological basis for the differences in retinal degeneration phenotype in the *Mertk* ^-/- V1^ *versus Mertk* ^-/- V2^ and *Mertk* ^-/- V3^ mouse lines can be trivially explained by a redundancy in phagocytosis provided for by TYRO3 in the absence of MERTK. This paradigm, nonetheless, failed entirely in explaining the remarkable anti-tumor resistance of *Mertk* ^-/- V1^ mice. *Tyro3* is not expressed in BMDMs, was not a DEG in BMDM RNAseq experiments and *Mertk* ^-/- V1^, *Mertk* ^-/- V2^ and *Mertk* ^-/- V3^ BMDMs were equally deficient in efferocytosis in *ex vivo* assay. Thus, neither TYRO3 nor any other DEG provided a redundancy to macrophage phagocytosis in the absence of MERTK. This observation seriously challenges the dogma that failure of MERTK-dependent efferocytosis of tumor cells universally improves anti-tumor immunity. Again, we do not propose to throw away the proverbial baby with the bath water. We only assert that anti-tumor immunity is contextual. Some indeed are dependent on disabling MERTK function in macrophages. For instance, macrophage-specific ablation of *Mertk* in B6 background conferred statistically significant reduction in tumor growth in the PyMT mouse model ^27^. That said, our results unequivocally point to independent modifiers of *Mertk* ^-/- V1^ mouse traits and calls for re-examination of the molecular and functional basis of phenotypes assigned to the loss of MERTK beyond anti-tumor immunity. An independent study showed that conditional ablation of *Mertk* in microglia led to decreased phagocytosis index in the mouse hippocampus ^5^. Paradoxically, hippocampal neurogenesis was impaired in *Mertk* ^-/- V1^ mice ^6^ or transiently increased in the absence of *Mertk* ^5^, suggesting laboratory-derived strain specific differences in *Mertk* ^-/- V1^ mice.

We cannot also entirely eliminate the possibility that *Mertk* itself does not have a functional role in some of the traits observed in *Mertk* ^-/- V1^ mice. For example, the DC-intrinsic defect in emigration to inflamed tissue that was initially ascribed to loss of function in *Nlrp10* in *Nlrp10* ^-/-^ mice first generated in a B6-BALB/c mixed background was subsequently revealed to be caused by a spontaneous mutation in *Dock8*, an unexpected byproduct of BALB/c variant of the gene ^36 37^. Similarly, *Il10* ^-/-^ mice in a B6 background differed in open-field behavioral tests from *Il10* ^-/-^ mice in a 129-B6 mixed background ^38^. The behavioral differences were found to be due to eQTLs *Emo4* and *Reb1* inherited from flanking region of *Il10* in chromosome 1 and not attributable to *Il10* itself ^39^.

In conclusion, two newly generated mouse lines with genetic ablation of *Mertk* reveal that two of the well-established *Mertk* ^-/- V1^ mice traits tested herein were both products of epistatic interactions with modifiers in the 129P2 genome. These new *Mertk* knockout mouse lines should prove useful for validation of phenotypes ascribed exclusively to *Mertk* based on studies employing *Mertk* ^-/- V1^ mice. Furthermore, these studies also identify a unique anti-tumor resistance in *Mertk* ^-/- V1^ mice against anti-CTLA-4 and anti-PD-1 refractory YUMM1.7 melanoma as well as against GL261 brain tumors, the molecular basis of which remains unknown. This model may yet prove useful for the discovery of a novel host anti-tumor response that can be therapeutically harnessed for improving outcomes in cancer.

## ACKNOWLEDGEMENTS

The authors would like to acknowledge the members of the Rothlin-Ghosh laboratory for scientific discussions relating to the preparation of this manuscript. This study was funded by NIH R01CA212376 (CVR and SG), NIH R01EY026215 grant (SCF), Howard Hughes Medical Institute Faculty Scholar award (CVR). SCF is supported by the Kim B. and Stephen E. Bepler Professorship in Biology. LDH was awarded NSF DGE-1122492 and Richard K. Gershon Fellowship.

## Materials and Methods

### Animals

Animals were bred and maintained under a strict 12-hour light cycle and fed with standard chow diet in a specific pathogen-free facility at Yale University. All animal experiments were performed in accordance with regulatory guidelines and standards set by the Institutional Animal Care and Use Committee of Yale University. All C57BL/6 mice were purchased from Jackson laboratories and subsequently bred and housed at Yale University. The widely used *Mertk* ^-/-V1^ mice have been described previously ^1,2^. *Mertk* ^-/-V1^ mice were crossed onto the C57BL/6J background (Jackson laboratories strain #: 000664) for at least ten generations. As shown in Figure 1, *Mertk* ^-/- V1^ mice have deletion of exon 17 that corresponds to the kinase domain of *Mertk* and the neomycin cassette is still present in the *Mertk* locus. Of note, exon 17 was referred to as exon 18 in the original description of the mouse ^2^.

To generate an independent line of mice that has *Mertk* deleted globally (referred to as *Mertk* ^-/-V2^ mice), we bred *Mertk* ^f/f^ mice with commercially available Rosa26^ERT2^Cre^+^ (Jackson laboratories strain #: 008463) mice. A description detailing the generation of the *Mertk* ^f/f^ mice can be found in Fourgeaud *et al*., 2016 ^3^. *Mertk* ^f/f^ mice originated from embryos of C57BL/6NJ background. Germline inactivation of the *Mertk* ^f/f^ allele in *Mertk* ^f/f^ Rosa26^ERT2^Cre^+^ mice was achieved by intraperitoneally injecting 3mg of 4-hydroxytamoxifen (Sigma-Aldrich) for five consecutive days. Two weeks post-tamoxifen injection, adult males and females were set up as breeder pairs. Litters from this cross were then screened for excision of the *Mertk* ^f/f^ allele. Once identified, excision positive mice were bred with C57BL/6J mice to eliminate the Cre recombinase. Finally, mice were genotyped to set apart *Mertk* ^-/-V2^ founder mice that had excised exon 18, encoding for the kinase domain of *Mertk*, on both alleles. Genome wide single nucleotide polymorphism (SNP) analysis of our established *Mertk* ^-/-V2^ mice indicated that ∼84.37% of their genome was C57BL/6J-derived while ∼15.62% was C57BL/6NJ-derived. An independent line targeting deletion of *Mertk*, designated as *Mertk* ^-/-V3^ mice, was generated at Cyagen Biosciences Inc. (Santa Clara, CA), by CRISPR/Cas9-mediated genome engineering. As shown in Figure 1, *Mertk* ^-/-V3^ mice have exons 3 and 4 targeted; transcribed mRNA from targeted allele with frameshift mutation undergo nonsense-mediated decay. Single guide RNAs (sgRNAs) were injected into fertilized C57BL/6NJ eggs and founder animals were identified by PCR followed by DNA sequencing analysis. Genome wide SNP analysis revealed that ∼55.22% of their genome was C57BL/6J-derived and ∼44.73% was C57BL/6NJ-derived.

CRISPR/Cas9 technique was used to make *Mertk* ^-/-V2^ *Tyro3* ^-/-V2^ mice, as described previously ^4^. In brief, T7-sgRNA templates were prepared by PCR, incorporating the guide sequences from the desired target regions in the mouse *Tyro3* gene (NCBI Gene ID: 22174), with a 5’ guide sequence of CTACACCTACAGAGAACAAG (sense orientation, cutting the gene at 8387/8 bp within intron 6) and a 3’ guide sequence of CCCAAGTGTCAGAATCCCAG (sense orientation, cutting the gene at 17800/1 bp within intron 18), thus resulting in a deletion of 9413 bp. The T7-sgRNA PCR templates were then used for *in vitro* transcription and purification with the MEGAshortscript T7 Transcription Kit and MEGAclear Transcription Clean-Up Kit, respectively (both from Thermo Fisher Scientific). Cas9 mRNA (CleanCap, 5-methoxyuridine-modified) was purchased from TriLink Biotechnologies. Subsequently, cytoplasmic microinjections of sgRNAs and Cas9 mRNA into single-cell embryos obtained from *Mertk* ^-/-V2^ donors were performed. Founder mice with heterozygous deletion of the *Tyro3* allele were identified with genotyping by PCR. Next, *Mertk* ^-/-V2^ *Tyro3* ^-/-V2^ line was established by heterozygote-to-heterozygote breeding. Consistent with previous reports^2,5^, male double knock- out mice lacking both *Mertk* and *Tyro3* have significantly smaller testicles and reduced fertility. *Tyro3* ^-/-V1^ mice have been described previously ^2^.

### PCR amplification

PCR reactions for sequencing were performed using primers listed in Table S1. PCR amplifications were carried out with TopTaq Master Mix Kit (Qiagen). PCR reactions of 25 μl were performed with 2 μl genomic DNA, 0.2 μM primer pair, 2.5 μl CoralLoad Concentrate 10x, 12.5 μl TopTaq Master Mix, 2x (contains TopTaq DNA polymerase, Toptaq PCR Buffer with 3 mM MgCl2 and 400 μM each dNTP). Thermal cycling conditions were based on Touchdown PCR method described by Korbie & Mattick, 2008 ^6^. PCR products were examined by gel electrophoresis.

### Short tandem repeat (STR) analysis

Genomic DNA was isolated from liver biopsies using the DNeasy blood and tissue kit (Qiagen) according to the manufacturer’s protocol. PCR reactions to amplify 24 different microsatellite regions on chromosome 2 were performed using primers listed in Table S1. Thermal cycling conditions were optimized based on Touchdown PCR method described by Korbie & Mattick, 2008 ^6^. Following PCR amplification, STR markers were scored by resolving PCR products on 4% agarose gels.

### Western blot

Protein lysates from adult mice RPE were obtained using a previously validated protocol ^7^. Briefly, the neural retina was removed and posterior eyecups were incubated on ice in 200μl of RIPA buffer with protease inhibitor cocktail, EDTA-free (Sigma-Aldrich, Inc.) for up to 1 hour. Posterior eyecups were removed and the dislodged RPE cells were sonicated for 20 seconds on ice. After 10 minutes at 14,000 rpm in a refrigerated centrifuge, supernatants were transferred to new tubes and protein content was quantified with Pierce BCA assay (Thermo Fisher Scientific) as per manufacturer’s instructions. Concomitantly, spleen, brain, testes and peritoneal exudate were collected from adult mice. Samples were kept in NP-40 buffer containing a cocktail of protease inhibitors (Sigma-Aldrich, Inc.). Tissues were mechanically disrupted and left rotating at 4°C for 2 hours to ensure complete homogenization. Subsequently, all samples were centrifuged for 20 min at 12,000 rpm and the supernatants was collected for protein quantification as described above.

For immunoblots, equal amounts of total protein in Laemmli Buffer were subjected to electrophoresis on precast polyacrylamide gels and transferred to PVDF membranes (Bio-Rad). Membranes were blocked and probed overnight with corresponding primary antibodies (MERTK: Abcam ab95925 1/1000, TYRO3: Cell Signaling Technology 5585S 1/1000, β-ACTIN: Cell Signaling Technology 8457S and 3700S). Secondary antibodies conjugated to near infrared fluorophores were detected using Odyssey Classic Imaging System (LI-COR Biosciences) and quantified with Image Studio Lite Software (LI-COR Biosciences).

### Generation of bone-marrow-derived macrophages (BMDMs)

BMDMs from age-matched, adult C57BL/6 *Mertk* ^-/-V1^, *Mertk* ^-/-V2^ and *Mertk* ^-/-V3^ mice were differentiated from bone marrow precursors. Briefly, bone marrow cells were isolated and propagated for 7 days in 30% L929-conditioned RPMI (Gibco) containing 20% FCS (Sigma-Aldrich) and 1% Pen/Strep (Gibco). At day 7, BMDMs were lifted for downstream assays.

### Phagocytosis assays

Thymocytes were isolated from 3-6-week-old C57BL/6 mice. Thymocytes were incubated for 4 hours with 1μg/ml of dexamethasone (Sigma-Aldrich) to induce apoptosis. In parallel, BMDMs from the indicated mice were collected and re-plated to adhere for 3 hours at 37°C. Apoptotic thymocytes, pre-labeled with 0.1mg/ml of pHrodo-SE (Thermo Fisher), were subsequently added to BMDMs at 6:1 ratio. Cells were co-cultured for 1 hour at 37°C. Afterwards, the apoptotic thymocyte-containing media was removed and BMDMs were washed 5 times with 1xPBS. Adherent BMDMs were then treated with Accutase (Sigma-Aldrich) for 10 minutes at 37°C. BMDMs were then gently scraped off for collection and stained for analysis with flow cytometry.

### Flow cytometry staining and acquisition

Single cell suspensions of BMDMs were stained in PBS with fixable viability dye (ThermoFisher Scientific) for 10 minutes. Cells were then incubated in 2% FCS/PBS solution containing anti-mouse CD16/32 antibody (BioLegend clone 93) for 15 minutes and subsequently stained with a combination of fluorophore-conjugated primary antibodies against mouse CD45 (BioLegend clone 30-F11), CD11b (BioLegend clone M1/70) and F4/80 (BioLegend clone BM8) at 4°C for 25 minutes. After staining, cells were washed and data was immediately acquired with BD LSRII flow cytometer using BD FACSDiva software (BD Biosciences). Finally, raw data were analyzed using FlowJo software (Tree Star Inc).

### Histological analysis

After mice were sacrificed by carbon dioxide inhalation, eyecups were immediately collected and incubated overnight in eye fixative (ServiceBio). Hematoxylin and eosin staining was performed by iHisto Inc. Samples were processed, embedded in paraffin, and sectioned at 4 μm. Paraffin sections were then deparaffinized and hydrated using the following steps: Xylene, two rounds,15 minutes each; 100% ethanol, two rounds, 5 minutes each; 75% ethanol, one round, 5 minutes; and 1x PBS, three rounds, 5 minutes each at room temperature. After deparaffinization, 4-μm-sectioned samples were placed on glass slides and stained with hematoxylin and eosin. Whole slide scanning (20x) was performed on an EasyScan Infinity (Motic). Outer nuclear layer (ONL) thickness was analyzed in ImageJ (NIH). Quantification of ONL thickness was performed in the medial retina. The areas analyzed were defined by distance from the optic disk. 10 measurements of ONL thickness were done per mouse.

### Electron microscopy

Adult mice were perfused with 1x PBS followed by 4% paraformaldehyde in PBS. Eyeballs were then carefully dissected, and further fixed in 2.5% glutaraldehyde and 2% paraformaldehyde in 0.1 M sodium cacodylate buffer (pH 7.4) for 1 hour at room temperature. Next, eyeballs were post-fixed in 1% OsO4 for 1 hour at room temperature and *en bloc* stained with 2% aqueous uranyl acetate for 30 minutes. They were then dehydrated in a graded series of ethanol, going from 70% to 100%, and finally transferred to 100% propylene oxide before being embedded in EMbed 812 resin, polymerized at 60°C overnight. Samples from medial retinas were cut respectively into thin sections of 60 nm by a Leica ultramicrotome (UC7), placed on standard EM grids and stained with 2% uranyl acetate and lead citrate. Retinal samples were examined with a FEI Tecnai transmission electron microscope at 80 kV accelerating voltage, digital images were recorded with an Olympus Morada CCD camera and iTEM imaging software at the Yale Center for Cellular and Molecular Imaging (CCMI) Electron Microscopy Facility.

### Electroretinography (ERG) recordings

All experimental animals were adapted in a dark room for 12 hours prior to recordings. Animals were anesthetized under dim red illumination using a 100 mg/kg Ketamine and 10 mg/kg xylazine cocktail injected intraperitoneally and pupils were dilated by application of a 0.5% tropicamide eye drop (Sandoz) at least 15 minutes before recordings. The cornea was intermittently irrigated with balanced salt solution to maintain the baseline recording and prevent keratopathy. Scotopic electroretinograms were acquired with UTAS ERG System with a BigShot Ganzfeld Stimulator (LKC Technologies, Inc.). A needle reference electrode was placed under the skin of the back of the head, a ground electrode was attached subcutaneously to the tail, and a lens electrode was placed in contact with the central cornea. The scotopic response was recorded for different luminances (i.e., log2 -2.1, -0.6, 0.4 and 1.4 cd.s/m^2^) using EMWin software, following manufacturer’s instructions (LKC Technologies, Inc.). The a-wave was measured as the difference in amplitude between baseline recording and the trough of the negative deflection, and the b-wave amplitude was measured from the trough of the a-wave to the peak of the ERG.

### Tumor implantations

100,000 YUMM1.7 melanoma cells, resuspended in 50ul of sterile 1xPBS, were subcutaneously injected into shaved rear flank of six- to twelve-week-old male mice. Mice were monitored for tumor growth by measuring the length and width of tumor masses using a caliper. Tumor volumes were scored with the formula (A x B^2^) x 0.4, in which A is the largest and B is the shortest dimension. Each mouse was said to have reached the end of its tumor-free-survival when the largest dimension of its tumor was measured to be 5mm. Mice were sacrificed once tumor growth reached an endpoint cutoff of 1000 mm^3^.

Anesthetized eight-to-twelve-week-old male mice were placed in a stereotactic apparatus and an incision was made with a scalpel over the cranial midline. A burr hole was made 1mm lateral and 2mm anterior to the bregma. A needle containing a suspension of GL261-luciferase (Luc) cells was inserted to a depth of 3mm. After the needle is allowed to rest in the burr hole for 5 minutes, 10,000 GL261-Luc cells were infused over the course of 4 minutes. Once cells were injected, the needle was allowed to rest in the skull for 5 minutes before removal. Finally, the incision was closed with vetbond tissue adhesive and animals were administered the full course of post-operative analgesic drugs, according to regulatory guidelines and standards set by the Institutional Animal Care and Use Committee of Yale University. Intracranial tumor growth was monitored using the 3D image reconstruction feature of the IVIS Spectrum instrument. Mice received an intraperitoneal injection of 150mg/kg of luciferin (Goldbio) prior to imaging. Tumor-bearing mice were checked daily for clinical signs of sickness behavior and were euthanized when one or more of the following symptoms were present: hunching, decreased activity, head tilt, weight loss, seizures and failure to groom.

### Generation of bone-marrow-derived dendritic cells (BMDCs)

BMDCs were differentiated from with granulocyte macrophage colony-stimulating factor (GM-CSF) as previously described ^8^. Briefly, bone marrow progenitors were collected from the femurs of adult C57BL/6 WT mice and 10 × 10^6^ progenitor cells were cultured in RPMI media (Gibco) supplemented with 10% FBS (Sigma Aldrich), 1% Penicillin-streptomycin (Gibco) and 20ng/ml of recombinant GM-CSF (Peprotech). On days 3 and 5, more supplemented media was added and cells were left in culture until day 7 when they would be ready to be used for generation of dendritic cell (DC) vaccines, as described below.

### Preparation of dendritic cell vaccines

GL261 tumor cell lysates were made by subjecting cells to six rounds of rapid freeze-thaw cycles, comprised of 3 minutes of incubation in liquid nitrogen and 4 minutes of incubation at 56°C. BMDCs were then incubated with GL261 tumor lysate (1mg of lysate/10×10^6^ BMDCs) for 2 hours at 37°C. After 2 hours, 1ug/ml LPS (Sigma) was added to BMDC-tumor lysate suspension and cells were incubated for 24 hours at 37°C. Next, supernatant was aspirated, BMDCs were collected and washed with 1xPBS three times. Finally, 1×10^6^ GL261-lysate pulsed BMDCs were intraperitoneally injected into mice at days 14 and 21 post intracranial implantation of GL261-Luc cells, as detailed above.

### Total RNA isolation and sequencing analysis

RNA was collected from postnatal day 25 mice using a previously validated method ^9^. Briefly, after euthanasia, mice eyes were enucleated and the posterior eyecup was incubated on ice in 400μl of RNAprotect (Qiagen) for 1 hour. RPE containing tubes were agitated for 10 minutes to dislodge any RPE cells attached to the posterior eyecup and centrifuged for 5 minutes at 685g. The RPE pellet was then subjected to total RNA extraction using RNeasy Mini kit (Qiagen) following manufacturer’s instructions. Similarly, total RNA from BMDMs was extracted from these cells using Rneasy Mini kit (Qiagen) following manufacturer’s instructions.

RNA libraries from BMDMs and RPE cells were prepared at the Yale Keck Biotechnology Resource Laboratory from three to six biological replicates per condition. Samples were sequenced using 150bp base pair paired-end reading on a NovaSeq 6000 instrument (Illumina). The raw reads were then subjected to trimming by btrim ^10^ to remove sequencing adaptors and low-quality regions. Next, reads were mapped to the mouse genome (GRCm38) using STAR ^11^. Finally, the Deseq2 ^12^ package was run to identify differentially expressed genes according to p-values adjusted for multiple comparisons. Genes with p-adjusted values less than 0.05 and log2 fold change ≤-1.25 or ≥1.25 were considered differentially expressed.

### Quantitative PCR analysis

Reverse transcription of RNA was performed utilizing iScript cDNA Synthesis Kit (Bio-Rad). Using KAPA SYBR Fast qPCR Kit (Kapa Biosystems), we amplified cDNA fragments and proceeded with qPCR reactions on CFX96 Thermal Cycler Real Time System (Bio-Rad). The reactions were normalized to 3 housekeeping genes (*Gapdh, Hprt* and *Rn18s*) and specificity of the amplified products was verified by looking at the dissociation curves. All oligonucleotides for qPCR were either purchased from Sigma-Aldrich or produced at the Yale University Keck Oligonucleotide Synthesis Facility (see sequences in Table S1).

### Statistical analysis

All statistical analyses were done using GraphPad Prism (GraphPad Software Inc.). All data are shown as mean ± S.E.M. and each data point represents a unique animal. Statistical differences between experimental groups were determined by employing various tests— namely, Kaplan-Meier test, Mann-Whitney test, Student’s t-test, one-way and two-way ANOVAs. Additionally, hypergeometric distribution analysis was employed to determine which chromosomes are over-represented in the pool of DEGs associated with each genotype. The distribution of DEGs was said to be enriched on a chromosome when the probability of association with a chromosome had p value <0.05, compared to the number of DEGs that would be expected to map to each chromosome by chance.

