## Supplementary Figure S1 for "Tissue-specific modifier alleles determine *Mertk* loss-of-function traits"

### Akalu *et al.* Supplementary Figure S1

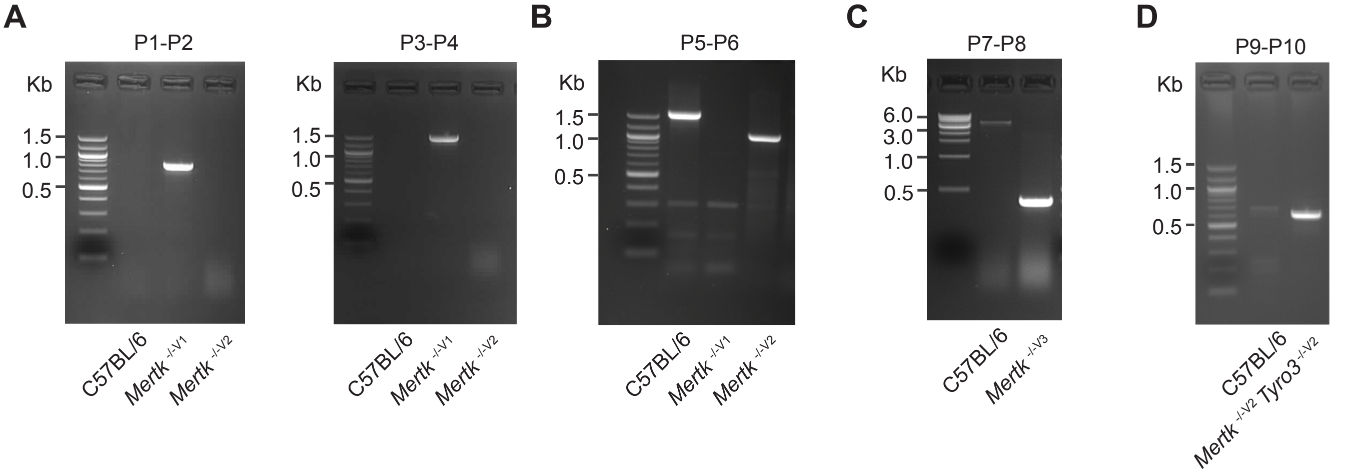


**Figure S1. PCR genotyping of mouse lines targeting *Mertk* (including a mouse line with the simultaneous ablation of *Mertk* and *Tyro3*).** (**A**) PCR amplification of genomic DNA (gDNA) from *Mertk*$^{-/-\,V1}$ mice using primers P1-P2 and P3-P4. Binding sites for primers P1-P2 and P3-P4 are indicated in **Figure 1B**. (**B**) Excision of targeted allele in *Mertk*$^{-/-\,V2}$ mice was confirmed using primers P5-P6. Binding sites for primers P5-P6 are indicated in **Figure 1B**. (**C**) PCR amplification of gDNA from *Mertk*$^{-/-\,V3}$ mice, using primers P7-P8, confirming excision. Binding sites for primers P7-8 are indicated in **Figure 1B**. (**D**) PCR amplification of genomic DNA from *Mertk*$^{-/-\,V2}$ *Tyro3*$^{-/-\,V2}$ mice, using primers P9-P10, confirming excision. Binding sites for primers P9-10 are indicated in **Figure 5C**.
