## Supplementary Figure S2 for "Tissue-specific modifier alleles determine *Mertk* loss-of-function traits"

### Akalu *et al.* Supplementary Figure S2

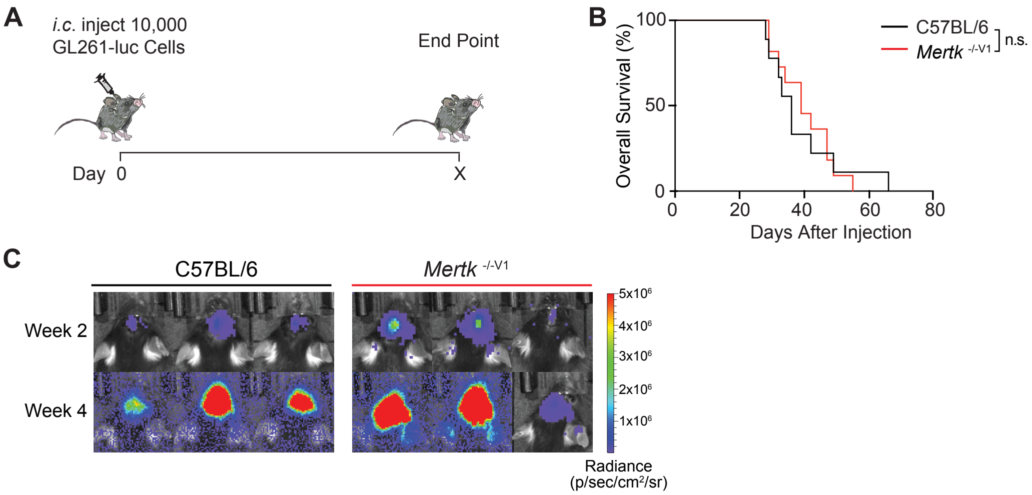


**Figure S2. Unvaccinated *Mertk* $^{-/-\ V1}$ mice are not protected against GL261 tumors.** (**A**) Schematic of intracranial injection of 10,000 GL261-Luc glioma cells in mice. (**B**) OS in C57BL/6 (n=9) and *Mertk* $^{-/-\ V1}$ (n=11) mice. n.s., Log-rank Mantel-Cox test. (**C**) Representative IVIS images of intracranial tumors in mice at D14 and D28 post-implantation.
