## Supplementary Figure S3 for "Tissue-specific modifier alleles determine *Mertk* loss-of-function traits"

### Akalu *et al.* Supplementary Figure S3

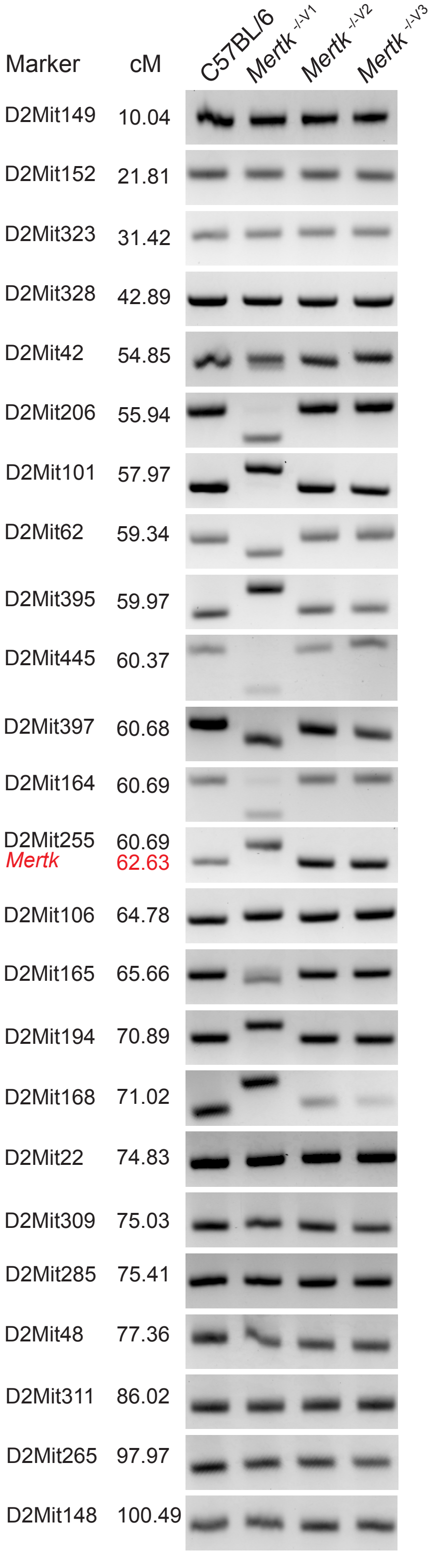


**Figure S3. A 129-specific genomic interval is present in *Mertk*$^{-/-V1}$ but not in *Mertk*$^{-/-V2}$ and *Mertk*$^{-/-V3}$ mice.** Representative PCR amplification products corresponding to 24 different microsatellite markers on chromosome 2 using genomic DNA extracted from C57BL/6, *Mertk*$^{-/-V1}$, *Mertk*$^{-/-V2}$ and *Mertk*$^{-/-V3}$ mice. Genotype at each microsatellite location was tested in at least 2-3 animals/genotype.
