## Supplementary Figure S4 for "Tissue-specific modifier alleles determine *Mertk* loss-of-function traits"

### Akalu *et al.* Supplementary Figure S4

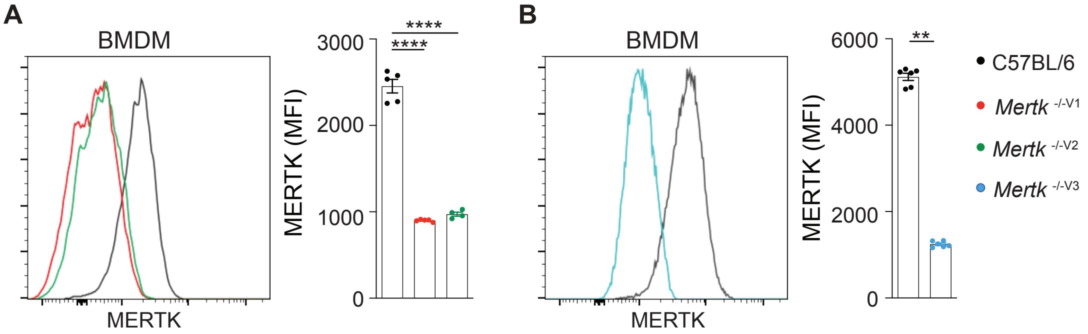


**Figure S4. Significantly diminished MERTK expression in BMDMs isolated from *Mertk* knockout mice.** (**A**, **B**) Representative flow-cytometry and independent quantification of MERTK levels in bone-marrow-derived macrophages (BMDMs) from C57BL/6, *Mertk* $^{-/- \, V1}$, *Mertk* $^{-/- \, V2}$ and *Mertk* $^{-/- \, V3}$ mice. (mean ± SEM of n=4-5 mice/genotype). ****p<0.0001, one-way ANOVA-Dunnet's test; **p<0.01, Mann-Whitney's test.
