## Supplementary Table S1 for "Tissue-specific modifier alleles determine *Mertk* loss-of-function traits"

| *Oligonucleotide* | *Sequence* |
|---|---|
| sgRNA#1 | GAAATTAATACGACTCACTATAGGGAGA**CTACACCTACAGAGAACAAG**GTTTTAGAGCTAGAAATAGCAAGTTAAAATAAGGCTAGTCCGTTATCAACTTGAAAAAGTGGCACCGAGTCGGTGCTTTTTT |
| sgRNA#2 | GAAATTAATACGACTCACTATAGGGAGA**CCCAAGTGTCAGAATCCCAG**GTTTTAGAGCTAGAAATAGCAAGTTAAAATAAGGCTAGTCCGTTATCAACTTGAAAAAGTGGCACCGAGTCGGTGCTTTTTT |
| Repair Oligo#1 | ATGAAGATCAATCACAGCTGATATTCCTCCCTCTTACATCCTGCTACTACACCTACAGAGAACAAGAGGAAGAGGAAGGCTAACAAACCCTGGCGAAGTTTGTCTGTCTGTCTGTTTGTTTGTTTG |
| Repair Oligo#2 | TTAGAGCTGAGAGGTACCCTAGACTTCAGATCCTCCTGCCACCACGCCCAAGTGTCAGAATCCCAGCGGTGTAGCACTTATGAGGTACTGGAGGTCATACCTGGGACTCTGTGCACACTGAGCAAG |
| P1 | GGGGCAGAGTACCTTGCTTT |
| P2 | CTGCGTGCAATCCATCTTGT |
| P3 | CTTTCGACCTGCAGCCAATATG |
| P4 | CCTCATCCCATATCAACACTGC |
| P5 | GCTCCAGCCCCTTTTACTTTTTGT |
| P6 | GATGTGCGATGTGATGGGAGGTAG |
| P7 | CTTAGAGACCAGGCAAGGTAGAAGCA |
| P8 | TCCTGAACACTCGCTGAATGCA |
| P9 | CAGGCCTGTGCTTTCTTTATGCTA |
| P10 | ACTGCTCTTCTGGGGGTTCTGA |
| *Mertk* F | GGCTTTTGGCGTGACCATGT |
| *Mertk* R | GGCCGTGGAGAAGGTAGTCG |
| *Tyro3* F | TGCCATTCGTACCGACTCAG |
| *Tyro3* R | TTTCTTGGACCCAGGACAGCTT |
| *Gchfr* F | CCACGCACCATGCCCTATCT |
| *Gchfr* R | GCTCCGGATCCGAGTGTTCA |
| *Gatm* F | TGCACTACATCGGCTCTCGG |
| *Gatm* R | CAGAGGATGGGTGGCCTTGT |
| *Thbs1* F | GGGGAGATAACGGTGTGTTTG |
| *Thbs1* R | CGGGGATCAGGTTGGCATT |
| *Pmepa1* F | AGCATGGAGATCACGGAGCTG |
| *Pmepa1* R | ACCGTACTCTCTGAGGGCCA |
| *Exd1* F | TCCCAGCAGTGACTACCATTT |
| *Exd1* R | CTCGTGCCCGAAGAACACTTT |
| *Itpka* F | CACGAAGCCGAGAGCAAGTG |
| *Itpka* R | TGAGCTGCCAATCACCTCGT |

| | |
|---|---|
| *Duoxa2* F | CGTTAACATTACACTCCGAGGAACA |
| *Duoxa2* R | CAGAATGCCACCCACAGTGT |
| *Aa467197* F | GGAGCCACATCTTTCGCTTTG |
| *Aa467197* R | CTCCTCAACGGGCTTCCATTG |
| *Gapdh* F | TCCCACTCTTCCACCTTCGA |
| *Gapdh* R | AGTTGGGATAGGGCCTCTCTT |
| *Hprt* F | AAGCTTGCTGGTGAAAAGGA |
| *Hprt* R | TTGCGCTCATCTTAGGCTTT |
| *Rn18s* F | GTAACCCGTTGAACCCCATT |
| *Rn18s* R | CCATCCAATCGGTACTAGCG |
| *D2Mit149* F | ATATCATATAGTAGAGAAAGCGTGCTG |
| *D2Mit149* R | TCATTAGACTTGGAAAAAAGTTTGC |
| *D2Mit152* F | CACAGATCTTGTAAGACCACGTG |
| *D2Mit152* R | TGCCATGAGTGTGGGACTAA |
| *D2Mit323* F | AGAATCCTAAGTGGTGGTTAGAGG |
| *D2Mit323* R | ACCCAAAGTTGTCTTTAAGTACACA |
| *D2Mit328* F | CTTTCAATGTTCCGGCATG |
| *D2Mit328* R | AAGACTTGCTTTCATTAGACCACA |
| *D2Mit42* F | ATTACTGGGCAGGAACATTTG |
| *D2Mit42* R | GCCAAACTTCCAGACTCCTC |
| *D2Mit206* F | TGTCAGAACTGGACAATGTCG |
| *D2Mit206* R | ATGATAACAGACACTAATGATTAGGGC |
| *D2Mit101* F | ATAATTCCTGATTTGCTGTTTGTG |
| *D2Mit101* R | ACATGAAGCCTAGAGGGTGC |
| *D2Mit62* F | GGATACCGTTTGGAAAGTAAACC |
| *D2Mit62* R | GCAAGAAGCACAGGAGGC |
| *D2Mit395* F | AGGTCAGCCTGGACTATATGG |
| *D2Mit395* R | AGCATCCATGGGATAATGGT |
| *D2Mit445* F | CCTATACACGCACACACAGACA |
| *D2Mit445* R | ATGCCCTGCTTGCTATTGTT |
| *D2Mit397* F | TGATGAAGGTTCTTTTTCTCCC |
| *D2Mit397* R | CCACAGTTGGTAATTATCTGGC |
| *D2Mit164* F | TCTCTGCTAATTAAGTTGAAGAGTGC |
| *D2Mit164* R | ACCAGTGTGTGTTTGTATGATGTG |
| *D2Mit255* F | GCAAGTGTGATCTGGGTGC |
| *D2Mit255* R | TGAGCACACTTACACTGTGGTG |
| *D2Mit106* F | GAGGGTTGCCAAAGAGACTG |
| *D2Mit106* R | CACCTCAGGGGAACATTGTG |
| *D2Mit165* F | TTTGGTCTTTCTAACCTTTGCA |
| *D2Mit165* R | AACAAAAACAAAACCAAAAAAACC |

| | |
|---|---|
| *D2Mit194* F | TGGAATTCCAAAGTCAAGGG |
| *D2Mit194* R | GGGAAGAATGGGGGAAGTTA |
| *D2Mit168* F | CTCACAGACACTGCACTATTACACA |
| *D2Mit168* R | TGTTCCTGCTATTGTTTTGGG |
| *D2Mit22* F | GCTCCCTTTCCTCTTGAACC |
| *D2Mit22* R | GGGCCCTTATTCTATCTCCC |
| *D2Mit309* F | ACAAATGCCACTCTCACATCC |
| *D2Mit309* R | TATTTCTCAGAGTCACTAGGAGTGATG |
| *D2Mit285* F | TCAATCCCTGTCTGTGGTAGG |
| *D2Mit285* R | TATGACACTTACAAGGTTTTTGGTG |
| *D2Mit48* F | GCTCTGCAGAAGATGCTGC |
| *D2Mit48* R | GCTGAGACGCAGAGTCGC |
| *D2Mit311* F | ACAGGCAGCCTTCCCTTC |
| *D2Mit311* R | TCTGTCCCGCTTCTGTTTCT |
| *D2Mit265* F | AATAATAATCAAGGTTGTCATTGAACC |
| *D2Mit265* R | TAGTCAAAATTCTTTTGTGTGTTGC |
| *D2Mit148* F | GTTCTCTGATCTACGGGCATG |
| *D2Mit148* R | TTCACTTCTACAAGTTCTACAAGTTCC |
